# Engineering Apical Integrin-binding Cellular Patches to Directed Cell Reprogramming via Mechanical Remodeling

**DOI:** 10.1101/2025.11.21.689608

**Authors:** Junchao Zhi, Tianrui Zhao, Wenjing Hou, Qizheng Zhang, Zhen Gao, Jiancan Yu, Kai Wu, Chenjie Xu, Xunwu Hu, Ye Zhang

## Abstract

Engineering extracellular microenvironments to control stem cell fate remains a central challenge in regenerative medicine. Here, we develop ECM-mimetic cellular patches formed by the supramolecular assembly of laminin-derived, integrin-binding ligands. The resulting fibrillar networks exhibit well-defined molecular packing and nanoscale ligand distribution, enabling specific engagement of apical integrin β1 on mesenchymal stem cells. This controlled interface converts molecular assembly into hierarchical mechanotransduction, coordinating cytoskeletal remodeling, nuclear deformation, and chromatin reorganization to drive neuronal reprogramming without genetic or chemical induction. Mechanistic studies reveal that the interplay between ligand assembly, spatial orientation, and network stability governs integrin activation and downstream transcriptional regulation These findings demonstrate how molecularly programmed assemblies can transform passive matrices into active, cell-instructive materials. This work establishes a framework for designing supramolecular systems that couple structural hierarchy with mechanotransductive control to direct stem cell fate and advance regenerative material strategies.

## Introduction

Neuronal replacement and regeneration are urgently needed for neurodegenerative diseases and traumatic injuries, yet the limited self-renewal of neurons in the adult central nervous system poses a major therapeutic challenge.^1^ While induced pluripotent stem cells (iPSCs)^2^ and neural stem cells (NSCs)^3^ offer routes for neuronal replacement,^4^ their clinical use is constrained by tumorigenicity,^5^ genetic instability,^6^ and limited accessibility.^7^ In contrast, mesenchymal stem cells (MSCs) are abundant, easily obtained, and compatible with autologous transplantation, making them attractive for regenerative medicine.^8^ Beyond their mesodermal potential, MSCs possess latent neurogenic plasticity that can be harnessed for neural repair.^9^ However, their robust cytoskeleton and mechanically reinforced nucleus predispose them toward stiffness-compatible fates such as osteogenesis, limiting neurogenic plasticity.^10, 11^ Conventional viral, chemical, or growth factor–based strategies^12, 13^ can transiently overcome these mechanical constraints but often yield unstable and heterogeneous phenotypes, with risks of genomic alteration or off-target signaling.^14^ Developing a non-genetic, mechanically defined strategy to unlock MSC neurogenic potential^15–17^ therefore remains a central challenge.

Cell fate is guided by both biochemical cues and mechanical signals, with the ECM serving as a critical interface for their integration.^18^ Among mechanical pathways, integrins act as primary sensors that transmit ECM-derived forces through the cytoskeleton to the nucleus,^19, 20^ reshaping nuclear architecture and epigenetic states to influence lineage specification.^21, 22^ Neuronal reprogramming, in particular, is favored in microenvironments with reduced mechanical signaling, such as soft substrates^23^ or attenuated cytoskeletal tension.^24^ Yet most current biomaterials focus on bulk stiffness or topographical features^25–29^ and provide limited means to modulate mechanotransduction at the membrane level with nanoscale precision, constraining spatially defined integrin engagement and coordinated force transmission. To address this, we engineered nanoscale cellular patches presenting an ECM-derived ligand capable of spatially engaging apical integrin β1. This defined activation orchestrates cytoskeletal tension, nuclear deformation, and chromatin remodeling, creating a permissive landscape for MSC neuronal reprogramming (Figure 1a). Unlike conventional approaches, this strategy enables non-genetic, exogenous-free control of lineage specification with high spatial and molecular precision, directly linking integrin engagement to cytoskeletal architecture, nuclear mechanics, and epigenetic state.

**Figure 1.**
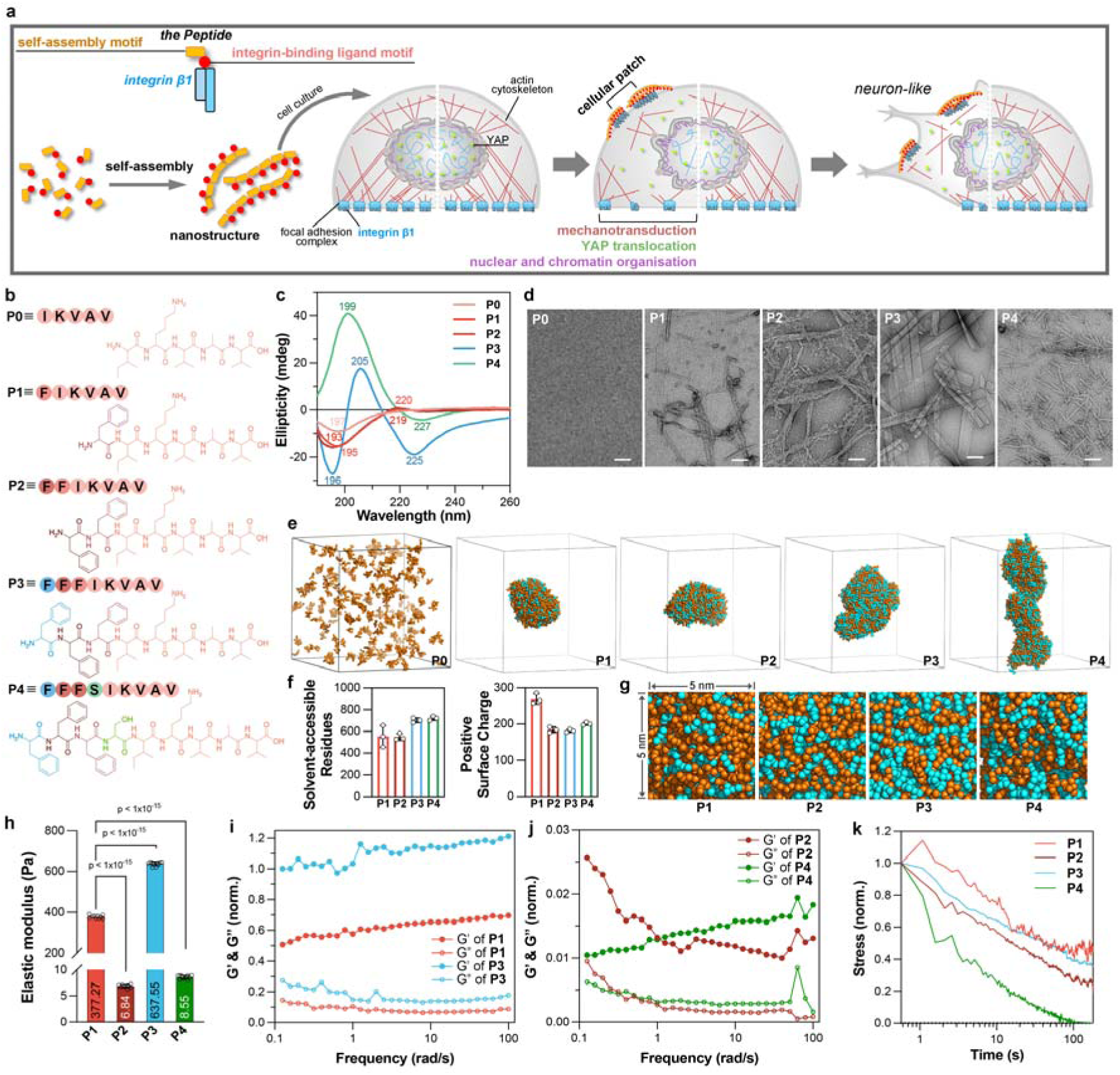
Engineering integrin-binding ligands for self-assembly into ECM-like nanostructures. **(a)** Schematic illustration of integrin-binding ligands self-assembling into a cellular patch that engages integrin clusters, initiates cytoskeletal reorganization, transmits mechanical cues to the nucleus, and ultimately induces MSC differentiation into neuron-like cells. **(b)** Chemical structures of the designed integrin-binding peptides. **(c)** CD spectra of peptides **P0**−**P4** in water at 200 μM. **(d)** TEM images of peptides **P0** (2 mM), **P1** (50 mM), **P2** (2 mM), **P3** (0.5 mM), and **P4** (0.5 mM) in water. Scale bars represent 100 nm. **(e)** Snapshots of simulated peptide self-assemblies. All simulations were conducted in a cubic box with a side length of 21.5 nm, corresponding to a peptide concentration of 50 mM. Phenylalanine residues are shown in cyan, and all other residues are shown in orange. Water molecules and ions are omitted for clarity. **(f)** Analysis of solvent-accessible residues and surface-exposed charge residues of simulated peptide assemblies (*n* = 3). **(g)** Representative snapshots of a 5 x 5 nm surface region of simulated peptide self-assemblies. Phenylalanine residues are shown in cyan, and all other residues in orange. **(h)** Elastic modulus of peptide assemblies in water (20 mg/mL) at room temperature (*n* = 10). Frequency sweep measurements of self-assemblies of **P1** and **P3 (i)**, **P2** and **P4 (j)**. **(k)** Stress relaxation behavior of peptide assemblies in water (20 mg/mL) at room temperature.

## Results

### Engineering Integrin-binding Ligand for Self-assembly

Building on our previously validated motif engineering strategy in which N-terminal aromatic extension promotes peptide self-assembly,^30, 31^ we designed a modular series of laminin-derived ligands in which biofunction and assembly are separable yet coupled. Specifically, the IKVAV motif,^32–34^ which represents a key basement membrane sequence that promotes cell adhesion and neurite outgrowth,^35^ was conjugated with one, two, or three L-phenylalanine residues at the N-terminus, generating assembling ligands **P1** (FIKVAV), **P2** (FFIKVAV), and **P3** (FFFIKVAV) (Figure 1b). Circular dichroism (CD) spectra revealed a gradual increase in ordered secondary structure, consistent with phenylalanine-driven supramolecular organization (Figure 1c). Transmission electron microscopy confirmed a corresponding morphological evolution. **P1** formed sparse and heterogeneous aggregates, **P2** produced thin nanofibers, and **P3** developed densely packed nanobelts (Figure 1d). To gain molecular-level insight, coarse-grained molecular dynamics simulations were performed (Figure 1e). Aggregation propensity and packing stability increased with aromatic valency, following the order **P3** > **P2** > **P1** (Figures S1–S3). These assemblies were driven mainly by hydrophobic and π–π interactions. In the highly cohesive **P3** assemblies, phenylalanine residues were buried in the hydrophobic core, while the IKVAV motif remained exposed on the surface. Energetic analyses indicated that hydrophobic and π–π interactions dominated the assembly formation and **P3** gained the greatest stability through efficient molecular packing (Figures 1f, S3, and S4). Ligand distribution snapshots revealed a transition from continuous to discontinuous presentation. Assemblies of **P1** maintained largely contiguous IKVAV surfaces, **P2** formed patchy domains, and **P3** exhibited isolated island-like distributions with enlarged gaps (Figure 1g). Such discontinuous nano-islands, coupled with high supramolecular cohesion, may generate mechanically stable and spatially organized adhesive interfaces that support robust integrin clustering.

To further probe how supramolecular packing and ligand topology influence interfacial presentation, we introduced a polar serine spacer between the FFF block and the IKVAV motif, generating **P4** (FFFSIKVAV). The spacer was designed to partially decouple the ligand from the aromatic core, attenuate overpacking, and increase interfacial ligand density while preserving bioactivity. This molecular configuration is consistent with the native laminin fragment A208,^33^ in which the SIKVAV submotif is flanked by polar residues that maintain surface exposure and support adhesion (Figure 1b). Circular dichroism spectra of **P4** displayed a distinct noncanonical pattern, and transmission electron microscopy revealed uniform thin nanofibers (Figures 1c and 1d), indicating a shift in supramolecular organization relative to **P3**. Coarse-grained simulations further confirmed a fine mosaic-like ligand distribution with elevated interfacial density (Figure 1g). Rheological analyses showed that **P3** formed the stiffest and most frequency-independent network, whereas **P4** exhibited faster stress relaxation, reflecting more dynamic molecular interactions (Figures 1h–k and S5). Together, these results indicate that systematic ligand engineering modulates supramolecular cohesion and nanoscale ligand topology, presentation, offering a tunable framework to explore how ECM-mimetic architectures regulate cellular responses.

### Ligand-assembled Cellular Patches Guide Neuronal Reprogramming of MSCs

Having established the supramolecular characteristics of the ligand series, we next evaluated their ability to form ECM-like structures and guide stem cell fate. Congo red staining revealed abundant assemblies on MSCs treated with **P3**, detectable signals with **P2** and **P4**, and negligible staining for **P0** and **P1** (Figure 2a and S6). SEM imaging confirmed ECM-like cellular patches adhering to MSC membranes, with morphology varying across the peptide series (Figure 2b and S7). **P1** produced sparse and small aggregates, while **P2** to **P4** generated fibrous networks that resembled their morphologies observed by TEM (Figure 1d). Neuronal induction was assessed using βIII-tubulin (TUBB3) immunostaining as an initial marker under transient or sustained treatments, with Blebbistatin serving as a positive control that reduces NMII-mediated contractility and phenocopies the pro-neurogenic effects of attenuated mechanotransduction.^36^ Sustained **P3** treatment at 200 μM yielded the highest efficiency of neuron-like cell induction, comparable to transient Blebbistatin at 20 μM or sustained Blebbistatin at 10 μM (Figure 2c and S8). Congo red staining further confirmed that sustained **P3** treatment preserved ECM-like cellular patches on the cell surface far more effectively than transient exposure (Figure 2d). Based on these results, sustained treatment at 200 μM was chosen for peptide screening. Live/dead assays confirmed that all peptide and Blebbistatin treatments were biocompatible (Figure S9). Using Nestin and TUBB3 as double markers, after 14 days of culture, **P3** induced neuron-like differentiation in more than 30% of MSCs, reaching an efficiency comparable to sustained Blebbistatin treatment. **P4** showed moderate effects, while **P0** to **P2** exhibited no significant activity (Figure 2e and 2f). qPCR analyses at both 7 and 14 days confirmed that **P3** significantly upregulated neuronal markers *TUBB3*, *MAP2*, and *NEFH* compared with all other treatments (Figure 2g and Table S1), while calcium-flux imaging revealed spontaneous calcium transients in more than 30% of **P3**-induced cells, consistent with acquisition of neuron-like excitability (Figure 2h and supporting video clip 1). Collectively, these findings reveal that the highly cohesive, discontinuous ligand topology of **P3**-assembled cellular patch is particularly suited for ECM-like assembly retention and mechanotransduction-mediated neuronal induction.

**Figure 2.**
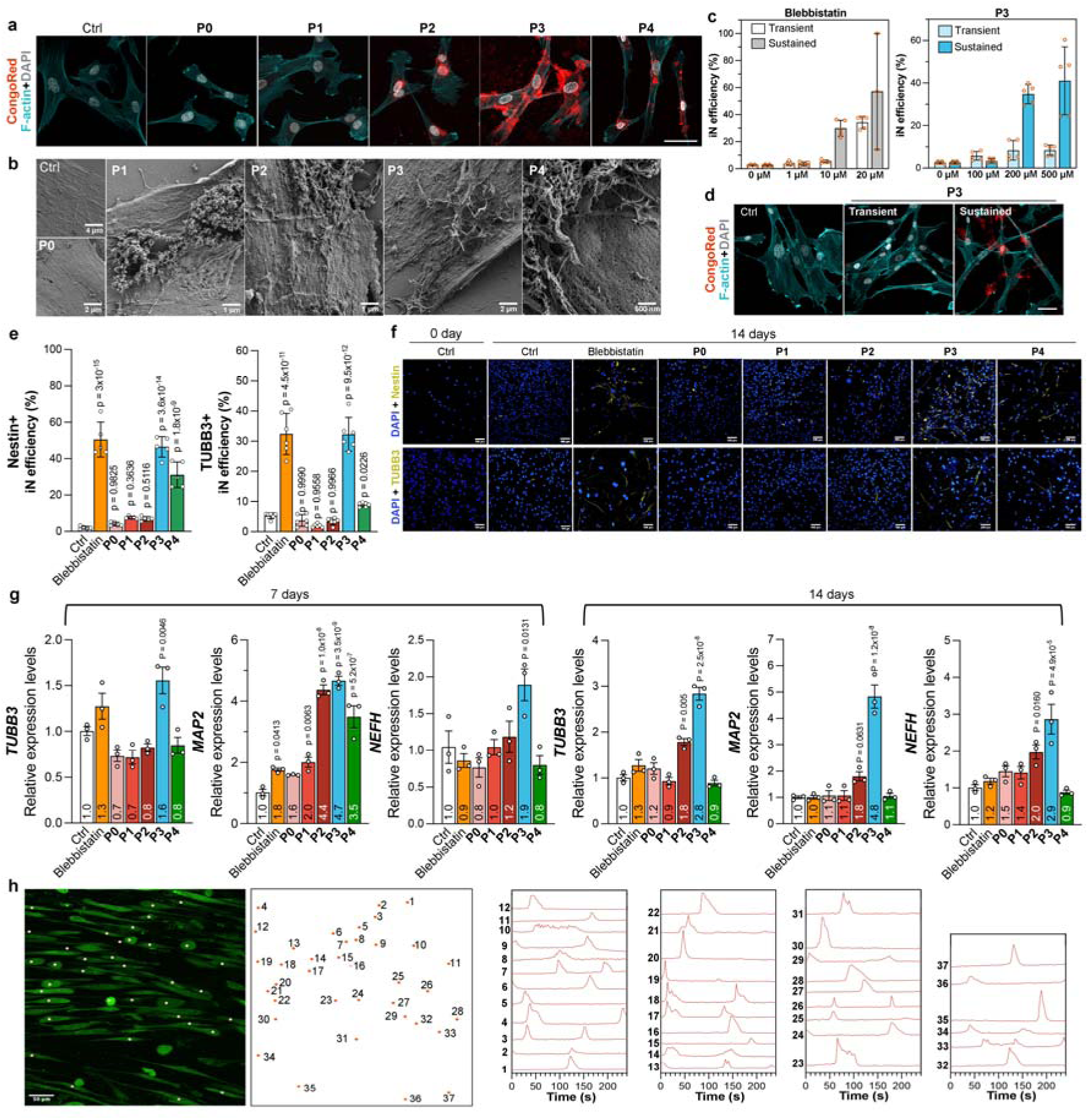
Peptide-assembled cellular patches instruct MSC differentiation into neuron-like cells. **(a)** Fluorescence images of MSCs with or without peptide treatment (200 μM, 1 day), co-stained with ActinGreen (cyan), DAPI (gray), and Congo Red (red). The scale bar represents 50 μm. **(b)** SEM images of MSCs with and without peptide treatment (200 μM, 1 day). **(c)** Reprogramming efficiency of MSCs with or without sustained Blebbistatin or **P3** treatment at different concentrations for 14 days, determined by TUBB3 staining (*n* = 5). **(d)** Fluorescence images of MSCs with or without peptide **P3** treatment (200 μM, transient or sustained for 14 days), co-stained with ActinGreen (cyan), DAPI (gray), and Congo Red (red). **(e)** Reprogramming efficiency of MSCs with or without sustained peptide treatment (200 μM, 14 days), determined by Nestin or TUBB3 staining (*n* = 5). Sustained treatment with Blebbistatin (10 μM) served as a positive control. **(f)** Representative immunofluorescence images corresponding to (e), with DAPI (blue), Nestin (yellow), and TUBB3 (yellow). The scale bars represent 100 μm. **(g)** Relative gene expression levels in MSCs following sustained treatment with Blebbistatin (10 μM) or peptides (200 μM) for 14 days, normalized to untreated MSCs (Ctrl) (*n* = 3). Fold-change values relative to Ctrl are indicated at the bottom of each bar. **(h) P3**-induced neuron-like cells labeled with Cal520. Yellow puncta mark cell positions corresponding to the red markers in the inset. Representative traces show spontaneous calcium transients. Statistical analysis was performed using one-way ANOVA compared with Ctrl. Data are presented as mean ± SD.

### Integrin β1 Engagement by Cellular Patches Remodels Focal Adhesion and Cytoskeleton Network

To verify the sequence specificity of ligand assemblies, a scrambled sequence of **P3** (**sP3**) was designed and characterized. Although **sP3** retained the ability to self-assemble (Figure 3b), its molecular packing differed from that of **P3** (Figure 3c), and Congo red staining revealed that it scarcely formed adherent cellular patches on MSC surfaces (Figure 3d). Consistently, **sP3** exhibited negligible neuronal induction efficiency (Figure 3e and 3f), underscoring the sequence-dependent effect. We next quantified integrin activation using an antibody recognizing its activated form.^37^ Confocal imaging revealed that **P3** treatment markedly increased integrin β1 activation (Figures 3g and 3h). Topographic mapping further showed that the enhanced signals were concentrated in the upper cellular layers, with no difference at the basal surface (Figure 3i). This spatial pattern indicates predominant integrin β1 activation on the apical membrane, consistent with the formation of apically localized cellular patches.

**Figure 3.**
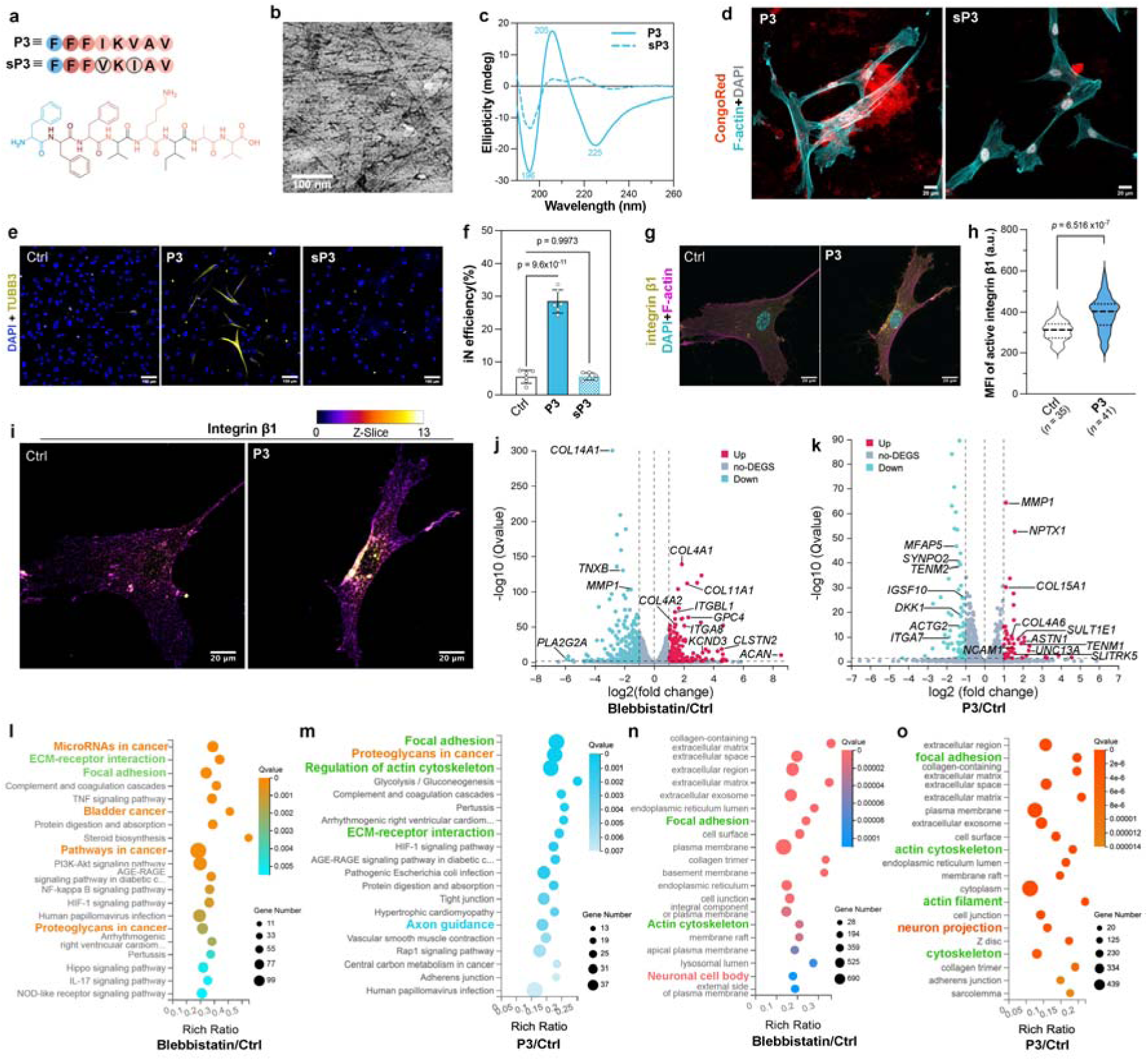
Cellular patch engages integrin β1 to remodel focal adhesion and actin cytoskeleton, driving MSC reprogramming. **(a)** Chemical structure of **P3**’s scrambled peptide **sP3**. **(b)** TEM image of **sP3** (1 mM) in water. **(c)** CD spectra of **P3** and **sP3** at 200 μM in water. **(d)** Fluorescence images of MSCs treated with **P3** or **sP3** (200 μM, 1 day), co-stained with ActinGreen (cyan), DAPI (gray), and Congo Red (red). The scale bars represent 20 μm. Representative immunofluorescence images **(e)** and quantification of reprogramming efficiency of MSCs **(f)** with or without sustained **P3** or **sP3** treatment (200 μM, 14 days), determined by TUBB3 staining (*n* = 5). The scale bars represent 100 μm. Representative images **(g)** and quantification of mean fluorescence intensity **(h)** of integrin β1 staining in MSCs upon treatment with **P3** co-stained with ActinGreen and DAPI compared with control on day 3. Scale bars, 20 μm. **(i)** Topographic map of z-positions of integrin β1 in MSCs corresponding to the images shown in (g). Scale bars represent 20 μm. Volcano plots of differentially expressed genes for MSCs treated with Blebbistatin **(j)** or **P3 (k)** versus control on day 14, with selected genes of interest highlighted. KEGG pathway enrichment analysis for MSCs upon sustained Blebbistatin **(l)** or **P3 (m)** treatment versus control on day 14. Go cellular component enrichment for MSCs upon sustained Blebbistatin **(n)** or **P3 (o)** treatment versus control on day 14. Statistical analysis was performed using one-way ANOVA compared with Ctrl. Data are presented as mean ± SD.

RNA sequencing revealed distinct transcriptional programs between Blebbistatin and **P3** treatments. Blebbistatin suppressed extracellular matrix remodeling genes such as *PLA2G2A*, *COL14A1*, *TNXB*, and *MMP1*, while upregulating collagens, adhesion related genes including *COL4A1*, *COL11A1*, *COL4A2*, *ITGBL1*, and neuronal priming markers *KCND3*, *CLSTN2*, and *ACAN*, suggesting stabilized ECM deposition and mild neurogenic bias (Figure 3j). In contrast, **P3** induced a divergent regulatory pattern characterized by downregulating of structural and regulatory genes such as *MFAP5*, *TENM2*, and *ITGA7*, accompanied by strong upregulation of neuronal adhesion and synaptic genes *NCAM1*, *SLITRK5*, and *NPTX1,*^38^ as well as matrix remodeling genes *MMP1* and *COL15A1* (Figure 3k). KEGG analysis revealed that Blebbistatin primarily enriched ECM–receptor interaction and focal adhesion, along with multiple cancer-related gene sets (e.g., microRNAs in cancer, pathways in cancer, bladder cancer, and proteoglycans in cancer) (Figure 3l). In contrast, **P3** selectively enriched focal adhesion, ECM–receptor interaction, actin cytoskeleton regulation, and axon guidance, indicating coordinated adhesion remodeling and neuronal signaling (Figure 3m). Consistent with this GO cellular component analysis identified **P3**-specific enrichment of actin filaments and neuron projections, signifying structural reorganization linked to neuronal differentiation (Figure 3n and 3o). Collectively, these findings underscore that ligand assembly–mediated apical integrin engagement elicits a physiological adhesion–cytoskeleton response that promotes neuronal reprogramming.

### Cytoskeletal Remodeling Downstream of Integrin Engagement Activates Mechanotransduction and Neuronal Reprogramming

Given that cellular patch-mediated integrin β1 activation occurred primarily at the apical membrane (Figure 3i), we examined how this spatial engagement influences cytoskeletal architecture and downstream mechanotransduction. At the initiation phase (day 3), both Blebbistatin and **P3** treatments reduced MSC spreading area (Figure 4a). By day 7, Blebbistatin-treated cells regained area, whereas **P3**-treated cells remained smaller, indicating sustained morphological constraint. Both treatments increased aspect ratio and decreased circularity, indicative of cell elongation and polarization, with **P3** eliciting a stronger effect. Perimeter analysis showed Blebbistatin reduced cell boundary complexity, while **P3** preserved peripheral features despite elongation. Paxillin staining revealed smaller focal adhesions in both groups (Figure 4b and 4c), accompanied by reduced phosphorylated myosin light chain (pMLC) intensity (Figure 4b and 4d). These changes indicated diminished actomyosin contractility and impaired adhesion maturation under reduced mechanical tension. F-actin organization diverged between treatments (figure 4e and 4f). Blebbistatin decreased both total and mean fluorescence intensity, consistent with its inhibition of myosin II ATPase activity that disrupt filament bundling. In contrast, **P3** treatment reduced total but increased mean intensity, reflecting an aligned and tension-bearing cytoskeletal network shaped by apical integrin engagement. In line with the adhesion remodeling visualized in cells, immunoblotting showed coordinated decreases in total and phosphorylated FAK and paxilline (Figures 4g and S10), indicating broad attenuation of integrin-dependent adhesion signaling. YAP subcellular localization showed decreased nuclear-to-cytoplasmic ratio under both treatments (Figures 4h and 4i), indicative of nuclear YAP export. **P3** induced the most pronounced cytoplasmic retention, aligning with enhanced neuronal lineage commitment. Overall, apical integrin β1 engagement by cellular patches initiates cytoskeletal and adhesion remodeling, sustains mechanotransduction, and directs MSC reprogramming along a neuronal trajectory.

**Figure 4.**
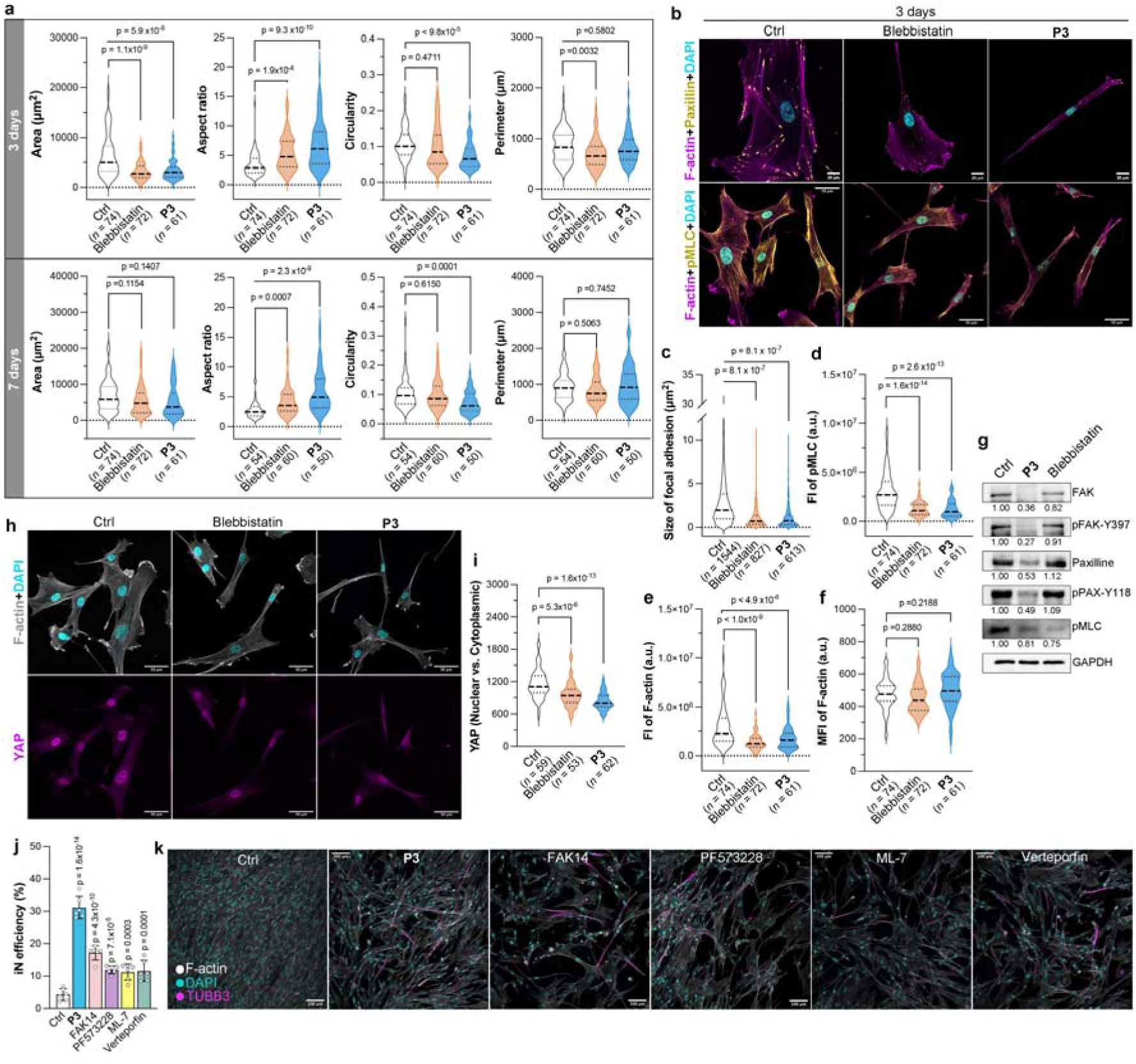
Cellular patch induces cytoskeletal remodeling to activate mechanotransduction driving MSC reprogramming. **(a)** Quantification of area, aspect ratio, circularity, and perimeter of MSCs upon sustained treatment with Blebbistatin or **P3** compared with control on day 3 and day 7. **(b)** Representative images of paxillin and pMLC staining in MSCs upon treatment with Blebbistatin or **P3** compared with control on day 3, costained with DAPI and F-actin. Scale bares, 20 μm for paxillin staining and 50 μm for pMLC staining. Quantification of paxillin size **(c)**, fluorescence intensity of pMLC **(d)**, fluorescence intensity of F-actin **(e)**, and mean fluorescence intensity of F-actin **(f)** in MSCs upon treatment with Blebbistatin or **P3** compared with control on day 3. **(g)** Immunoblotting analysis of FAK, pFAK-Y397, paxillin, pPAX-Y118, and pMLC expression in MSCs treated with Blebbistatin and **P3**. GAPDH serves as a loading control. **(h)** Representative images of YAP staining in MSCs upon treatment with Blebbistatin or **P3** compare with control on day 3. Scale bars 50 μm. **(i)** Quantification of the ration of YAP intensities in the nucleus relative to that in the cytoplasm of MSCs upon the treatment with Blebbistatin or **P3** compared with control on day 3. Quantification of reprogramming efficiency **(j)** and the representative immunofluorescence images **(k)** of MSCs treated with **P3,** FAK14, PF573228, ML-7, and Verteporfin treatment, determined by TUBB3 staining (*n* = 5). The scale bars represent 100 μm. Statistical analysis was performed using one-way ANOVA compared with Ctrl. Data are presented as mean ± SD.

To delineate the contribution of individual mechanotransduction components, pharmacological inhibitors targeting distinct mechanosensitive nodes were employed. FAK14 and PF573228 were used to inhibit FAK-mediated focal adhesion signaling, ML-7 to block MLCK-dependent actomyosin contractility, and Verteporfin to suppress YAP/TAZ–TEAD transcriptional activity. After 14 days of treatment, quantification of TUBB3-positive cells revealed enhanced neuronal induction cross all inhibitor-treated groups relative to control, with the efficiency ranking **P3** > FAK14 > PF573228 ≈ ML-7 ≈ Verteporfin (Figures 4j and 4k). FAK14 produced a stronger effect than PF573228, consisitent with its partial inhibition of FAK that preserves residual scaffolding functions required for limited adhesion and cytoskeletal signaling. In contrast, PF573228 more completely suppresses FAK activity, placing it on par with ML-7 and Verteporfin, whose inhibition of actomyosin contractility or YAP nuclear signaling similarly attenuates neuronal induction. Collectively, these results establish that **P3**-mediated integrin β1 engagement orchestrates hierarchical cytoskeletal remodeling, activating a coordinated mechanotransduction cascade spanning focal adhesion dynamics, actomyosin contractility, and YAP nuclear signaling. This integrated and spatially coordinated mechanotransduction-mediated regulation surpasses the effect of perturbing individual downstream nodes, highlighting the advantage of upstream integrin engagement in directing MSC reprogramming toward neuronal fate.

### Cellular Patch Remodels Nuclear Architecture and Histone Modification to Support MSC Reprogramming

Since YAP export reflects reduced cytoskeletal tension, we next examined whether this mechanical change shift propagates to the nucleus.^21, 39^ Lamin A/C staining^40–42^ revealed that **P3** treatment markedly enhanced nuclear mean fluorescence intensity, expanded nuclear area, and increased both the nuclear aspect ratio and wrinkle index, indicating pronounced nuclear elongation and mechanical deformation (Figure 5a and 5b). These features parallel previously reported nuclear responses to altered adhesion mechanics and cytoskeleton tension.^22, 43, 44^ In contrast, Blebbistatin induced a larger nuclear area but increased circularity with minimal changes in aspect ratio or wrinkle index, reflecting nuclear relaxation rather than mechanically driven deformation. These distinctions indicate that **P3**-generated cellular patches uniquely reinforce lamin A/C organization and impose directed nuclear remodeling, whereas global actomyosin inhibition lacks this spatial mechanotransduction input.

**Figure 5.**
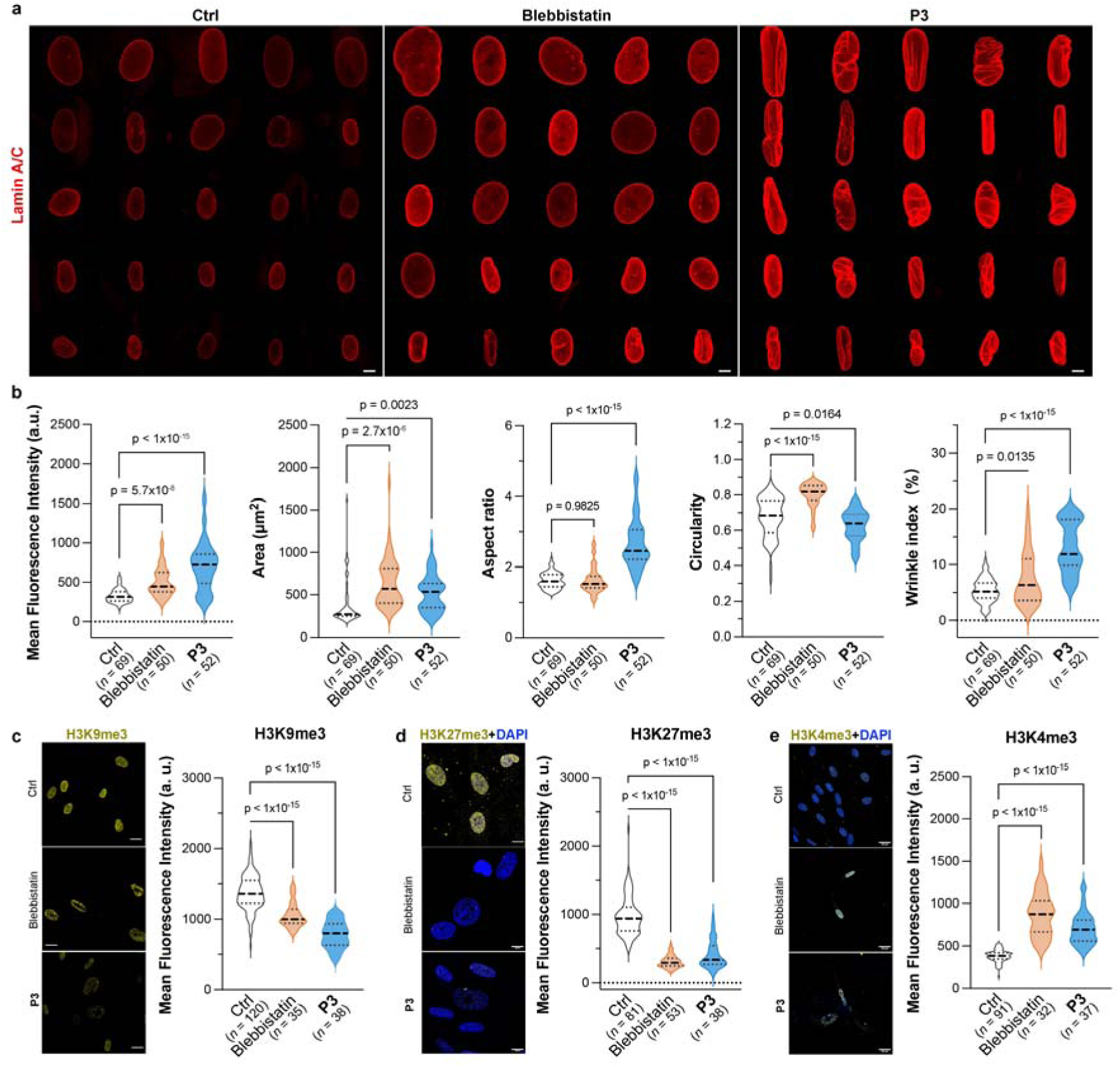
Lamin A/C-mediated epigenetic reorganization underlies cellular patch-induced MSC reprogramming. **(a)** Representative images of lamin A/C staining in MSCs upon sustained treatment with Blebbistatin or **P3** compared with control on day 14. Scale bars, 10 μm. **(b)** Quantification of nuclear mean fluorescence intensity, area, aspect ratio, circularity, and wrinkle index of MSCs upon sustained treatment with Blebbistatin or **P3** compared with control on day 14, coresponding to the imaging results in (a). Representative images of H3K9me3 staining and the quantification of H3K9me3 mean intensity **(c-e)** Representative images and quantification of histone modification in MSCs upon sustained treatment with Blebbistatin or **P3** compared with control on day 14, including H3K27me3 (c, scale bars 20 μm ), H3K27me3 (d, scale bars 10 μm), and H3K4me3 (e, scale bars 20μm). Statistical analysis was performed using one-way ANOVA compared with control.

The nuclear structural alterations prompted us to access whether ask whether **P3**-mediated deformation also affects chromatin state. **P3** treatment substantially reduced H3K9me3 (Figure 5c) and H3K27me3 (Figure 5d), reflecting relaxation of constitutive and facultative heterochromatin. Concurrently, H3K4me3, associated with active transcription, was increased, indicating enhanced transcriptional permissiveness (Figure 5e). Blebbistatin produced milder and less coordinated effects. H3K9me3 decreased modestly, H3K27me3 reduction was similar to **P3**, and H3K4me3 showed a slight increase, reflecting generalized tension loss rather than targeted mechanotransduction. Together, these findings indicate that sustained apical integrin β1 engagement by **P3** cellular patches drives a hierarchical mechanotransduction program that links cytoskeletal remodeling to nuclear architecture and selective epigenetic reorganization.^45, 46^ By reinforcing lamin A/C, deforming the nucleus, and modulating histone marks, **P3** establishes a permissive chromatin landscape that primes MSCs for neuronal reprogramming, highlighting the advantage of spatially controlled upstream mechanotransduction over global actomyosin inhibition.

## Discussion

This work demonstrates how supramolecular organization and ligand orientation can be harnessed to create predictable, translatable, and programmable extracellular environments. By leveraging a basal membrane-derived motif (IKVAV) applied apically, we introduce a striking contrast in orientation that reveals new opportunities for materials-guided mechanotransduction. This inversion, achieved by repurposing a native basal signal for apical engagement, enables precise orchestration of cytoskeletal tension, nuclear deformation, and chromatin remodeling, illustrating how molecular-level ligand design and supramolecular assembly can exert hierarchical control over cell behavior.

The resulting ECM-mimetic cellular patches act as active, instructive surfaces rather than passive scaffolds, bridging the gap between conventional stiffness- or topography-based biomaterials and fully programmable, cell-instructive platforms. By controlling nanoscale ligand distribution, network viscoelasticity, and motif accessibility, these assemblies translate molecular design into functional regulation of integrin engagement and downstream mechanotransduction. This counterintuitive strategy provides a generalizable blueprint for engineering ECM-mimetic materials that actively guide cellular fate, integrating molecular precision with functional instruction to pave the way for next-generation biomaterials.

## Methods

### Peptide synthesis

Peptides **P0**, **P1**, **P2**, **P3**, and **P4** were custom-synthesized by solid-phase synthesis with a purity of ≥ 95% (GL Biochem, China). The structure of **P3** was previously reported and validated by NMR and LC-MS in our earlier work. Characterization data for the remaining peptides are provided below.

### P0: IKVAV

^1^H NMR (600 MHz, DMSO-*d*_6_): δ 8.44 (d, *J* = 7.6 Hz, 1H), 8.08 (s, 2H), 8.02 – 7.93 (m, 2H), 7.84 (d, *J* = 8.6 Hz, 1H), 7.78 (s, 2H), 4.42 – 4.30 (m, 2H), 4.15 (dd, *J* = 8.4, 6.9 Hz, 1H), 4.12 – 4.08 (m, 1H), 3.63 (s, 1H), 2.72 (dd, *J* = 13.2, 7.3 Hz, 2H), 2.46 (dt, *J* = 3.6, 1.9 Hz, 1H), 2.06 – 1.89 (m, 2H), 1.79 – 1.71 (m, 1H), 1.64 – 1.56 (m, 1H), 1.50 (dt, *J* = 20.2, 6.2 Hz, 3H), 1.34 (d, *J* = 7.6 Hz, 1H), 1.29 – 1.21 (m, 1H), 1.20 – 1.12 (m, 3H), 1.12 – 1.03 (m, 1H), 0.87 – 0.75 (m, 18H) ppm. LCMS (ESI): for C_25_H_48_N_6_O_6_, [M+H]^+^ calcd 529.37, found 529.5 (*m*/*z*).

### P1: FIKVAV

^1^H NMR (600 MHz, DMSO-*d*_6_): δ 8.58 (d, *J* = 8.6 Hz, 1H), 8.21 (d, *J* = 7.8 Hz, 1H), 8.14 (s, 2H), 8.03 (d, *J* = 7.3 Hz, 1H), 7.87 (d, *J* = 8.6 Hz, 1H), 7.82 (s, 1H), 7.79 (d, *J* = 8.9 Hz, 2H), 7.32 – 7.25 (m, 3H), 7.25 – 7.21 (m, 2H), 4.38 (p, *J* = 7.1 Hz, 1H), 4.32 (dd, *J* = 14.0, 8.1 Hz, 1H), 4.26 (t, *J* = 8.0 Hz, 1H), 4.18 (dd, *J* = 8.8, 6.5 Hz, 1H), 4.14 (dd, *J* = 8.6, 5.6 Hz, 2H), 3.06 (dd, *J* = 14.6, 4.6 Hz, 1H), 2.92 (dd, *J* = 14.1, 7.8 Hz, 1H), 2.75 (dd, *J* = 13.2, 7.4 Hz, 2H), 2.05 (dq, *J* = 13.6, 6.8 Hz, 1H), 1.96 (dd, *J* = 13.4, 6.7 Hz, 1H), 1.74 – 1.68 (m, 1H), 1.68 – 1.62 (m, 1H), 1.59 – 1.52 (m, 2H), 1.52 – 1.44 (m, 2H), 1.39 – 1.25 (m, 2H), 1.19 (d, *J* = 7.1 Hz, 3H), 1.11 – 1.06 (m, 1H), 0.88 – 0.78 (m, 18H) ppm. HRMS (ESI): for C_34_H_57_N_7_O_7_, [M+H]^+^ calcd 676.4393, found 676.4405 (*m*/*z*).

### P2: FFIKVAV

^1^H NMR (600 MHz, DMSO-*d*_6_) δ 8.75 (d, *J* = 8.1 Hz, 1H), 8.29 (d, *J* = 8.5 Hz, 1H), 8.10 (d, *J* = 8.0 Hz, 1H), 8.03 (d, *J* = 7.6 Hz, 3H), 7.87 (d, *J* = 8.4 Hz, 1H), 7.77 (d, *J* = 9.0 Hz, 3H), 7.31 – 7.23 (m, 9H), 7.20 (t, *J* = 7.1 Hz, 1H), 4.72 (td, *J* = 8.9, 4.2 Hz, 1H), 4.38 (p, *J* = 7.4 Hz, 1H), 4.33 (td, *J* = 8.1, 5.5 Hz, 1H), 4.26 (t, *J* = 8.2 Hz, 1H), 4.19 (dd, *J* = 8.9, 6.2 Hz, 1H), 4.14 (dd, *J* = 8.4, 5.5 Hz, 1H), 4.04 – 3.98 (m, 1H), 3.11 (dd, *J* = 14.3, 4.5 Hz, 1H), 3.01 (dd, *J* = 14.2, 4.1 Hz, 1H), 2.93 (dd, *J* = 14.3, 7.9 Hz, 1H), 2.80 (dd, *J* = 14.3, 9.7 Hz, 1H), 2.75 (dd, *J* = 15.6, 6.2 Hz, 2H), 2.05 (dq, *J* = 13.2, 6.8 Hz, 1H), 1.96 (h, *J* = 6.4 Hz, 1H), 1.75 (ddt, *J* = 9.9, 6.9, 3.3 Hz, 1H), 1.65 (dq, *J* = 16.0, 6.0 Hz, 1H), 1.58 – 1.42 (m, 4H), 1.38 – 1.25 (m, 2H), 1.19 (d, *J* = 6.9 Hz, 3H), 1.15 – 1.06 (m, 1H), 0.89 – 0.78 (m, 18H) ppm. HRMS (ESI): for C_43_H_66_N_8_O_8_, [M+H]^+^ calcd 823.5077, found 823.5052 (*m*/*z*).

### P4: FFFSIKVAV

^1^H NMR (500 MHz, DMSO-*d*_6_) δ 12.56 (s, 1H), 8.64 (d, *J* = 8.0 Hz, 1H), 8.40 (d, *J* = 8.1 Hz, 1H), 8.25 (d, *J* = 7.6 Hz, 1H), 8.07 (d, *J* = 7.9 Hz, 1H), 8.04 – 7.90 (m, 3H), 7.86 (d, *J* = 8.6 Hz, 1H), 7.80 (d, *J* = 8.5 Hz, 1H), 7.72 (d, *J* = 8.8 Hz, 1H), 7.70 – 7.58 (m, 2H), 7.32 – 7.13 (m, 15H), 5.04 (s, 1H), 4.70 (td, *J* = 8.8, 4.2 Hz, 1H), 4.61 (td, *J* = 8.6, 4.5 Hz, 1H), 4.42 (dd, *J* = 13.2, 5.9 Hz, 1H), 4.40 – 4.34 (m, 1H), 4.30 (dd, *J* = 14.4, 8.8 Hz, 1H), 4.28 – 4.24 (m, 1H), 4.17 (dd, *J* = 8.4, 6.8 Hz, 1H), 4.14 (dd, *J* = 8.5, 5.7 Hz, 1H), 3.96 (dd, *J* = 7.4, 4.3 Hz, 1H), 3.62 (d, *J* = 4.4 Hz, 1H), 3.57 (d, *J* = 5.0 Hz, 1H), 3.09 – 3.00 (m, 3H), 2.87 – 2.79 (m, 2H), 2.79 – 2.71 (m, 3H), 2.05 (dq, *J* = 13.4, 6.7 Hz, 1H), 1.96 (dq, *J* = 13.4, 6.6 Hz, 1H), 1.80 – 1.71 (m, 1H), 1.69 – 1.60 (m, 1H), 1.56 – 1.46 (m, 3H), 1.45 – 1.37 (m, 1H), 1.36 – 1.23 (m, 2H), 1.20 (d, *J* =.0 Hz, 3H), 1.15 – 1.04 (m, 1H), 0.87 (d, *J* = 6.8 Hz, 6H), 0.85 – 0.82 (m, 6H), 0.82 – 0.78 (m, 6H) ppm. LCMS (ESI): for C_55_H_80_N_10_O_11_, [M+H]^+^ calcd 1057.61, found 1057.50 (*m*/*z*).

### Transmission Electron Microscopy (TEM)

Carbon-coated copper grids were glow-discharged to enhance surface hydrophilicity. A 10 μL aliquot of the sample solution was applied to each grid and incubated for 10 s before excess liquid was blotted off with filter paper. Grids were rinsed three times with 5 μL of distilled water, air-dried briefly, and stained with 5 μL of 1% (w/v) uranyl acetate (SPI supplies, USA; 02624-AB) for 10 s. Excess stain was removed, and the grids were air-dried for an additional 10 s. The morphology of the molecular assemblies was examined under high vacuum using a JEM-F200 transmission electron microscope (JEOL, Japan).

### Simulation Procedures

Peptide structures were predicted using AlphaFold3 and converted to MARTINI coarse-grained (CG) models with martinize2 in the Vermouth framework, defining coiled-coil helices as the secondary structure. Four FIKVAV-derived peptides (FIKVAV, FFIKVAV, FFFIKVAV, and FFFSIKVAV) carried a net charge of +1 due to a single protonated lysine, with no further protonation adjustments. Systems containing 300 peptides (50 mM, 3.46–5.63 wt%) were solvated in a 21.5 nm cubic box with MARTINI water and counterions. After energy minimization (≤ 5000 steps; max force < 100 pN), self-assembly was simulated for 2.5 µs in the NPT ensemble (303 K, 1 bar) using GROMACS 2025.2 with a 25 fs timestep. Nonbonded interactions used a 1.1 nm cutoff, electrostatics employed the reaction-field method (εr = 15), and pressure control used the stochastic cell rescaling barostat. Peptide aggregation was assessed by clustering within a 0.6 nm cutoff, and structural fluctuations were quantified as RMSF over the final 1 µs of three equilibrated trajectories. RMSF values were mapped to normalized B-factors for visualization using VMD.

### Mechanical characterization of peptide assemblies

Peptide assemblies were prepared by dissolving peptide powders in Milli-Q water at a concentration of 20 mg/mL. The pH of each solution was adjusted to 7.0 using 1M NaOH to initiate gelation. At this concentration, all four peptides formed self-assembled hydrogels.

Rheological measurements were performed on a controlled-stress rheometer (HR-30, TA Instruments) equipped with a 20 mm diameter parallel-plate geometry. All tests were conducted at 25°C, with a constant gap of 100 μm. For each measurement, 100 μL of hydrogel was placed on the Peltier plate using a calibrated micropipette. The upper plate was gently lowered to the set gap, and excess material was trimmed with a laboratory spatula to ensure uniform contact and a consistent meniscus.

The viscoelastic behavior of the peptide hydrogels was characterized using a multi-step protocol. Strain amplitude sweeps were first conducted at a fixed angular frequency of 6 rad/s over a strain range of 0.01%−300% to define the linear viscoelastic region (LVR). Frequency sweeps were then performed within the LVR across 0.01−100 rad/s to obtain the storage modulus (G’) and loss modulus (G’’). Young’s modulus values were estimated from the frequency-dependent mechanical data. Stress relaxation tests were conducted by applying a constant strain within the LVR and monitoring stress decay over 180 s to assess network stability.

### Cell Culture of hBMSCs

Human bone marrow-derived mesenchymal stem cells (hBMSC) at passage 2 were obtained from Oricell (China; catalog no. HUXMF-01001). Cells were expanded in 75 cm^2^ tissue culture-treated flasks using Bone Mesenchymal Stem Cell Basal Medium (MBSCBM; Oricell, China) and subcultured twice. Cells at passage 4 were harvested using TrypLE™ Express (Gibco, USA; 2993993) and used for all subsequent experiments.

### Confocal microscopy

Confocal imaging was performed using an AX R Confocal Microscope System (Nikon, Japan). To assess the F-actin organization, cells were seeded on glass-bottom culture dishes (801002, NEST) and treated with peptides as indicated. After treatment, the medium was removed, and the cells were washed with pre-warmed PBS. A freshly prepared solution of Congo Red (0.1 mg/mL; Abcam, UK; ab145645, Lot GR3426274-1) in culture medium was added, and the cells were incubated for 30 min at 37 °C. Cells were then washed tree times with DPBS and fixed with 4% paraformaldehyde in phosphate buffer solution (Beyotime, China; P0099) for 20 min at room temperature, followed by three additional DPBS washes. Nuclei and F-actin were stained with NucBlue^TM^ Fixed Cell Stain ReadyProbes^TM^ reagent (Invitrogen, USA; 2831256) and Actin-Green 488 (Invitrogen, USA; 2647610) for 15 min at room temperature. After a final DPBS wash, cells were imaged by confocal microscopy. Morphological features were quantified using ImageJ.

### Scanning Electron Microscopy (SEM)

After peptide treatment, the culture medium was removed, and cells were washed three times with DPBS. Cells were fixed with 2.5% glutaraldehyde in 0.1 M cacodylate buffer for 30 min, followed by post-fixation with 1% OsO_4_ (SPI supples, USA; 02604-AB) in 0.1 M cacodylate buffer (EMD Millipore, USA; 4012482) for 30 min. Samples were washed three times with Milli-Q water (5 min each), dehydrated through a graded ethanol series, rinsed with t-BuOH (Innochem, China; KDHFT01), and freeze-dried in a lyophilizer (VTLG-10C, Yetuo, China) for over 12 h. Before imaging, samples were coated with a 5 nm platinum layer using a platinum sputter coater (EM ACE600, Leica). SEM images were acquired using an ultra-high-resolution FE-SEM (Gemini 300, ZEISS, Germany) operated at 1.0 kV.

### Immunostaining

Cells were fixed with 4% paraformaldehyde for 30 min and blocked with Immunostaining Blocking Solution (Beyotime, China; P0260) for 1 h. Primary antibodies incubation was then performed using the following antibodies: anti-beta3-Tubulin (1: 200, 5568S, Cell Signaling), anti-Nestin (1: 200, 73349S, Cell Signaling), anti-Paxillin (1:50, ab32084, Abcam), anti-P-Myosin Light Chain2(1: 200, 3674S, Cell Signaling), anti-YAP (1: 50, ab52771, Abcam), anti-Lamin A/C (1: 200, 4777S, Cell Signaling), anti-H3K4me3 (1:200, ab8580, Abcam), anti-H3K9me3 (1:200, ab8898, Abcam), anti-H3K27me3 (1:200, ab6002, Abcam), diluted in 1% BSA for 1 h at 4 °C.

After washing three times with DPBS, cells were incubated with the appropriate secondary antibodies: Goat Anti-Mouse lgG H&L (Alexa Fluor® 568) (ab175473, Abcam, 1:1000), Goat Anti-Rabbit lgG H&L (Alexa Fluor® 568) (ab175471, Abcam, 1:1000). Goat Anti-Mouse lgG H&L (Alexa Fluor® 647) (ab150115, Abcam, 1:1000), Goat Anti-Rabbit lgG H&L (Alexa Fluor® 647) (ab150075, Abcam, 1:1000), diluted in 1% BSA for 1 h at room temperature in the dark.

When preparing cells for integrin imaging, cells were fixed in 4% paraformaldehyde for 30 min and blocked with 3% BSA (SRE0098, Sigma-Aldrich) for 30 min. Cells were then incubated with anti-Integrin β1 (MAB2079Z, Sigma-Aldrich) diluted in 1% BSA for 1 h at 4 °C. After three washes with PBS, cells were incubated with Goat Anti-Mouse IgG H&L (Alexa Fluor® 568) (1:1000, ab175473, Abcam) diluted in 1% BSA for 1 h at room temperature in the dark. Following another three washes with PBS, nuclei and F-actin were stained using NucBlue™ Fixed Cell Stain ReadyProbes™ reagent and ActinGreen™ 488 for 15 min at room temperature.

### RNA Isolation and Quantitative PCR (qPCR) Analysis

The expression levels of neurogenesis-related genes, including *MAP2* (Microtubule-Associated Protein 2), *TUBB3* (Tubulin Beta 3 Class III), and *NEFH* (Neurofilament Heavy Chain), were quantified by real-time PCR. Total RNA was extracted from cultured cells using TRIzol® reagent (Invitrogen, USA; 10091011) according to the manufacturer’s instructions. Complementary DNA (cDNA) was synthesized from 2 μg of purified total RNA using the High-Capacity cDNA Reverse Transcription Kit (Applied Biosystems, USA; 2824962). The resulting cDNA served as the template for qPCR amplification.

Each qPCR reaction (20 μL total volume) contained 10 μL of Power SYBR^TM^ Green PCR Master Mix (Applied Bio-systems, USA; A57156), 7.4 μL of nuclease-free water, 1.6 μL of cDNA, and 1 μL of primer mix. Amplification was performed on a CFX Connect^TM^ Real-Time PCR Detection System (Bio-Rad, USA) using 40 cycles of thermal cycling. Primers sequences used in this study are listed in Table S1 (Supporting Information).

### Calcium-Flux Imaging

Cells were loaded with the calcium indicator dye Cal-520 AM following standard procedures. A 2 mM stock solution of Cal-520 AM (AAT Bioquest, USA; 21130) was prepared in anhydrous dimethyl sulfoxide (DMSO). The working solution (4 µM) was obtained by diluting the stock solution in Hanks’ Balanced Salt Solution (HBSS; Gibco, USA; 2896482) containing calcium and magnesium, supplemented with 0.02% Pluronic F-127 (Invitrogen, USA; 2935433) to enhance dye dispersion. After removing the culture medium, cells were incubated with the Cal-520 AM working solution for 30-60 min at 37°C in a humidified incubator with 5% CO₂, protected from light. Following incubation, cells were washed three times (5 min each) with pre-warmed, dye-free HBSS (without Ca²⁺/Mg²⁺; Gibco, USA; 3245605) to remove excess extracellular dye. The cells were then allowed to recover for 30 min in fresh buffer to ensure complete de-esterification of the AM ester. Time lapse fluorescence imaging was performed using an AX R laser-scanning confocal microscope (Nikon, Japan).

### RNA-seq and Differential Gene Expression Analysis

The culture medium was removed, and adherent cells were gently rinsed with 1.5 mL of sterile DPBS (pH 7.2–7.4). After aspirating the buffer, TRIzol**^®^** Reagent was added directly to lyse the cells. The lysate was pipetted to ensure complete detachment and homogenization, then incubated at room temperature for 5 min. The resulting cell lysate was transferred to 1.5 mL microcentrifuge tubes, sealed with parafilm, and stored at −80 °C until shipment. Samples were transported on dry ice to BGI Genomics (Wuhan, China) for sequencing on the DNBSEQ platform. Differentially expressed genes (DEGs) were identified using the DEGseq2 pipeline, with significance thresholds set at q ≤ 0.05 and log_2_FC ≥ 1.0, where FC denotes fold change.

### Western Blotting Assay

For Western blotting, the following primary antibodies were used: Phospho-Myosin Light Chain 2 (Thr18/Ser19) (1:1000, 3674S, Cell Signaling Technology), Phospho-Paxillin (Tyr118) (1:1000, 69363S, Cell Signaling Technology), Paxillin (1:1000, ab32084, Abcam), Phospho-FAK (Y397) (1:1000, ab81298, Abcam), FAK (1:1000, 3285S, Cell Signaling Technology), and GAPDH (1:1000, ab8245, Abcam). HRP-conjugated Goat Anti-Mouse IgG (1:5000, G21040, Invitrogen) and Goat Anti-Rabbit IgG (1:5000, G21234, Invitrogen) were used as secondary antibodies.

Cells were lysed in RIPA buffer (Thermo Fisher Scientific) supplemented with protease and phosphatase inhibitors (Thermo Fisher Scientific) on ice for 30 min. Lysates were centrifuged at 12,000 × *g* for 15 min at 4 °C, and supernatants were collected. Protein concentration was measured using a BCA Protein Assay Kit (Thermo Fisher Scientific). Equal amounts of protein (20–50 μg per lane) were resolved by SDS–PAGE (8–12% gels) and transferred to PVDF membranes (Millipore) using the Trans-Blot® Turbo™ Transfer System (Bio-Rad) at 25 V for 30 min.

Membranes were blocked with 5% BSA in TBST (TBS, ST661, Beyotime + 0.1% Tween-20, P9416, Merck) for 1 h at room temperature and then incubated with primary antibodies overnight at 4 °C. After three washes with TBST, membranes were incubated with HRP-conjugated secondary antibodies for 1 h at room temperature. Protein bands were visualized using enhanced chemiluminescence (ECL, Thermo Fisher Scientific) and imaged with a ChemiDoc System (Bio-Rad). Band intensities were quantified using ImageJ (NIH), with protein levels normalized to GAPDH.

### Statistical Analysis

No statistical methods were used to predetermine sample size. The experiments were not randomized, and the investigators were not blinded to allocation during data collection or analysis. All measurements were performed on 1–3 independent biological replicates from separate experiments. The exact sample size (*n*) and statistical test used for each dataset are specified in the corresponding figure legends. Statistical analyses were carried out using GraphPad Prism (Version 10.6.1, GraphPad Software, USA). Data are presented as mean ± s.e.m. or s.d., as indicated in the figure legends. *P* values were adjusted for multiple comparisons where applicable, and exact *p* values are reported in the figures.Statistical significance was defined as *p* < 0.05.

## Acknowledgements

This work was supported by Songshan Lake Materials Laboratory, Guangdong S& T program (2024 A 0505040003), Guangdong Basic and Applied Basic Research Foundation (2025 A 1515010616 ), the Pearl River Talent Program of Guangdong Province (2023 CX10C027), SLAB Young Scientists Program, SLAB AI+Materials Program.

## Author contributions

X.H. and Y.Z.conceived the study and design the experiments. J.Z., T.Z., W.H., Q.Z., and Z.G. performed the experiments. J.Z. conducted circular dichroism spectroscopy, TEM and SEM sample preparation, live/dead assay, quantitative RT-PCR, confocal microscopy, calcium-flux imaging, westen blotting; T.Z. conducted mechanical characterizations; W.H. condicted simulations; Q.Z. conducted the NMR spectroscopy and MS spectroscopy; J.Z. and Z.G. conducted the sample preparation for RNA-seq; X.H., Y.Z., J.Y., K.W. and C.X. advised the study. X.H. and Y.Z. wrote the paper.

## Competing interests

The authors declare the following competing financial interest(s): X.H. and Y.Z. are inventors on a patent application related to the data presented in this manuscript.

## Data and materials availability

All data are available in the main text or the supplementary materials.

## References

1. Götz, M. & Bocchi, R. Neuronal replacement: Concepts, achievements, and call for caution. Curr Opin Neurobiol 69, 185–192 (2021).

2. Hu, B.Y. et al. Neural differentiation of human induced pluripotent stem cells follows developmental principles but with variable potency. Proc Natl Acad Sci U S A 107, 4335–4340 (2010).

3. Nie, L. et al. Directional induction of neural stem cells, a new therapy for neurodegenerative diseases and ischemic stroke. Cell Death Discovery 9, 215 (2023).

4. Zhou, B. et al. Effects of Univariate Stiffness and Degradation of DNA Hydrogels on the Transcriptomics of Neural Progenitor Cells. Journal of the American Chemical Society 145, 8954–8964 (2023).

5. Lee, A.S., Tang, C., Rao, M.S., Weissman, I.L. & Wu, J.C. Tumorigenicity as a clinical hurdle for pluripotent stem cell therapies. Nature Medicine 19, 998–1004 (2013).

6. Steinemann, D., Göhring, G. & Schlegelberger, B. Genetic instability of modified stem cells - a first step towards malignant transformation? Am J Stem Cells 2, 39–51 (2013).

7. Trounson, A. & McDonald, C. Stem Cell Therapies in Clinical Trials: Progress and Challenges. Cell Stem Cell 17, 11–22 (2015).

8. Han, X. et al. Mesenchymal stem cells in treating human diseases: molecular mechanisms and clinical studies. Signal Transduction and Targeted Therapy 10, 262 (2025).

9. Lau, D. et al. Stem Cell Clinics Online: The Direct-to-Consumer Portrayal of Stem Cell Medicine. Cell Stem Cell 3, 591–594 (2008).

10. Mathieu, P.S. & Loboa, E.G. Cytoskeletal and Focal Adhesion Influences on Mesenchymal Stem Cell Shape, Mechanical Properties, and Differentiation Down Osteogenic, Adipogenic, and Chondrogenic Pathways. Tissue Engineering Part B: Reviews 18, 436–444 (2012).

11. Oliver-De La Cruz, J., et al. Substrate mechanics controls adipogenesis through YAP phosphorylation by dictating cell spreading. Biomaterials 205, 64–80 (2019).

12. Mitchell, A.C., Briquez, P.S., Hubbell, J.A. & Cochran, J.R. Engineering growth factors for regenerative medicine applications. Acta Biomaterialia 30, 1–12 (2016).

13. Hwang, K.C. et al. Chemicals that modulate stem cell differentiation. Proc Natl Acad Sci U S A 105, 7467–7471 (2008).

14. Menche, C. & Farin, H.F. Strategies for genetic manipulation of adult stem cell-derived organoids. Experimental & Molecular Medicine 53, 1483–1494 (2021).

15. Jia, X. et al. Adaptive Liquid Interfacially Assembled Protein Nanosheets for Guiding Mesenchymal Stem Cell Fate. Advanced Materials 32, 1905942 (2020).

16. Jia, X. et al. Modulation of Mesenchymal Stem Cells Mechanosensing at Fluid Interfaces by Tailored Self-Assembled Protein Monolayers. Small 15, 1804640 (2019).

17. Jia, X. et al. Adaptive liquid interfaces induce neuronal differentiation of mesenchymal stem cells through lipid raft assembly. Nat Commun 13, 3110 (2022).

18. Hussey, G.S., Dziki, J.L. & Badylak, S.F. Extracellular matrix-based materials for regenerative medicine. Nature Reviews Materials 3, 159–173 (2018).

19. Elosegui-Artola, A. et al. Rigidity sensing and adaptation through regulation of integrin types. Nat Mater 13, 631–637 (2014).

20. Kong, F. et al. Cyclic Mechanical Reinforcement of Integrin&#x2013;Ligand Interactions. Molecular Cell 49, 1060–1068 (2013).

21. Song, Y. et al. Biphasic regulation of epigenetic state by matrix stiffness during cell reprogramming. Science Advances 10, eadk0639.

22. Wu, Y. et al. Viscoelastic extracellular matrix enhances epigenetic remodeling and cellular plasticity. Nature Communications 16, 4054 (2025).

23. Teixeira, A.I. et al. The promotion of neuronal maturation on soft substrates. Biomaterials 30, 4567–4572 (2009).

24. Di, X. et al. Cellular mechanotransduction in health and diseases: from molecular mechanism to therapeutic targets. Signal Transduction and Targeted Therapy 8, 282 (2023).

25. Harati, J. et al. Defined Physicochemical Cues Steering Direct Neuronal Reprogramming on Colloidal Self-Assembled Patterns (cSAPs). ACS Nano 17, 1054–1067 (2023).

26. Jiang, X. et al. Nanofiber topography and sustained biochemical signaling enhance human mesenchymal stem cell neural commitment. Acta Biomaterialia 8, 1290–1302 (2012).

27. Poudineh, M. et al. Three-Dimensional Nanostructured Architectures Enable Efficient Neural Differentiation of Mesenchymal Stem Cells via Mechanotransduction. Nano Letters 18, 7188–7193 (2018).

28. Yang, L., Jurczak, K.M., Ge, L. & van Rijn, P. High-Throughput Screening and Hierarchical Topography-Mediated Neural Differentiation of Mesenchymal Stem Cells. Adv Healthc Mater 9, e2000117 (2020).

29. Yoo, J. et al. Nanogrooved substrate promotes direct lineage reprogramming of fibroblasts to functional induced dopaminergic neurons. Biomaterials 45, 36–45 (2015).

30. Chen, Y. et al. Formulate Adaptive Biphasic Scaffold via Sequential Protein-Instructed Peptide Co-Assembly. Advanced Science 11, 2401478 (2024).

31. Hu, X. et al. Control cell migration by engineering integrin ligand assembly. Nature Communications 13, 5002 (2022).

32. Barcellona, M.N. et al. Control of adhesive ligand density for modulation of nucleus pulposus cell phenotype. Biomaterials 250, 120057 (2020).

33. Kasai, S. et al. Multifunctional peptide fibrils for biomedical materials. Peptide Science 76, 27–33 (2004).

34. Nomizu, M. et al. Cell binding sequences in mouse laminin alpha1 chain. J Biol Chem 273, 32491–32499 (1998).

35. Yu, T.T. & Shoichet, M.S. Guided cell adhesion and outgrowth in peptide-modified channels for neural tissue engineering. Biomaterials 26, 1507–1514 (2005).

36. He, Z.-Q. et al. Pharmacological Perturbation of Mechanical Contractility Enables Robust Transdifferentiation of Human Fibroblasts into Neurons. Advanced Science 9, 2104682 (2022).

37. Álvarez, Z. et al. Bioactive scaffolds with enhanced supramolecular motion promote recovery from spinal cord injury. Science 374, 848–856 (2021).

38. Diener, C. et al. Paving the way to a neural fate – RNA signatures in naive and trans-differentiating mesenchymal stem cells. European Journal of Cell Biology 103, 151458 (2024).

39. Kim, D.H. & Wirtz, D. Cytoskeletal tension induces the polarized architecture of the nucleus. Biomaterials 48, 161–172 (2015).

40. Cosgrove, B.D. et al. Nuclear envelope wrinkling predicts mesenchymal progenitor cell mechano-response in 2D and 3D microenvironments. Biomaterials 270, 120662 (2021).

41. Dickinson, R.B. & Lele, T.P. Nuclear shapes are geometrically determined by the excess surface area of the nuclear lamina. Frontiers in Cell and Developmental Biology Volume 11–2023 (2023).

42. Kim, J.-K. et al. Nuclear lamin A/C harnesses the perinuclear apical actin cables to protect nuclear morphology. Nature Communications 8, 2123 (2017).

43. Song, Y. et al. Transient nuclear deformation primes epigenetic state and promotes cell reprogramming. Nat Mater 21, 1191–1199 (2022).

44. Ayushman, M. et al. Cell tumbling enhances stem cell differentiation in hydrogels via nuclear mechanotransduction. Nature Materials 24, 312–322 (2025).

45. Soto, J. et al. Reduction of Intracellular Tension and Cell Adhesion Promotes Open Chromatin Structure and Enhances Cell Reprogramming. Adv Sci (Weinh) 10, e2300152 (2023).

46. Song, Y. et al. Asymmetric Cell Division of Fibroblasts is An Early Deterministic Step to Generate Elite Cells during Cell Reprogramming. Advanced Science 8, 2003516 (2021).

