## Supplemental Text, table, figures for "Engineering Apical Integrin-binding Cellular Patches to Directed Cell Reprogramming via Mechanical Remodeling"

Supporting figures S1-S10

Supporting Table S1

Attachments 1-7

### Materials and methods

#### Materials

Uranyl acetate and osmuim tetroxide ($OsO_4$) were purchased from SPI Supplies (USA). Sodium hydroxide was obtained from Sigma-Aldrich (USA). Cacodylate buffer was sourced from EMD Millipore (USA), n-Butanol was purchased from Innochem (China). 2-Propanol was purchased from Sigma-Aldrich (USA). Anhydrous ethanol (purity ≥ 99.5%) was purchased from Nanjing Chemical Reagent Co., Ltd. (Nanjing, China). All reagents were of analytical grade and used without further purification.

#### Peptide synthesis

Peptides P0–P4 were custom-synthesized by solid-phase peptide synthesis (≥95% purity; GL Biochem, China). P3 was previously developed and characterized for a different study [1], where its structure was confirmed by NMR and LC–MS. Characterization data for the remaining peptides are provided.

#### Circular Dichroism (CD) Spectroscopy

Peptides were dissolved in Milli-Q water to prepare 10 mM stock solutions, with quantities calculated based on their molecular weights. The pH was adjusted to 7.0 using 1M sodium hydroxide (NaOH) under continuous vortexing to ensure homogeneous mixing. Working solutions at the desired concentrations were prepared by serial dilution in the same aqueous medium. To ensure complete molecular reorganization and formation of stable nanostructures, peptide solutions were incubated at room temperature for 12 h before CD measurement. CD spectra were acquired at room temperature on a JASCO J-1700 spectropolarimeter using a 1 mm path-length quartz cuvette. The bandwidth was set to 0.5 nm, and spectra were collected over a range of 185-280 nm.

**Simulation Procedures**

Peptide structures were first predicted using AlphaFold3[2] and subsequently converted into MARTINI coarse-grained (CG) models via martinize2 within the Vermouth framework[3].

To better capture the secondary structural characteristics of the sequences, input topology files were manually adjusted during CG mapping, with helical regions explicitly reassigned as β-sheet types.

All FIKVAV-derived peptides—namely FIKVAV, FFIKVAV, FFFIKVAV, and FFFSIKVAV—contain only one charged residue, the naturally protonated lysine (+1). Experimental reports have confirmed that these peptides can spontaneously self-assemble into fibrous aggregates.therefore, no protonation state modifications were applied during coarse-graining, and the default protonation states defined in the MARTINI force field were retained. As a result, each peptide carried a net charge of +1.

Simulations were performed in two stages within a cubic simulation box of 21.5 × 21.5 × 21.5 $nm^3$, solvated with MARTINI water and neutralized with counterions. In the initial setup, 300 peptide molecules were randomly distributed in the box with a minimum inter-peptide distance of 3Å, corresponding to a peptide concentration of approximately 50 mM and a weight fraction ranging from 3.46% to 5.63%, depending on sequence length. This concentration range is commonly employed to accelerate self-assembly dynamics in silico and is roughly fivefold higher than typical experimental conditions.

Energy minimization was carried out using the steepest descent algorithm for up to 5000 steps, or until the maximum force fell below 100 pN, with positional restraints of 2000 $kJ \cdot mol^{-1} \cdot nm^{-2}$ applied to backbone beads. A cutoff of 1.1 nm was used for all nonbonded interactions. Electrostatic interactions were treated using the reaction-field method with a relative dielectric constant of 15. The Lennard–Jones potential was handled with a potential-shift scheme as defined in the MARTINI force field [4].

Subsequent CG-MD simulations were performed using GROMACS 2025.2 [5] in the NPT ensemble with a 25 fs integration time step[7]. Temperature was maintained at 303 K using the V-rescale thermostat ($\tau_T$ = 1 ps) [4], and pressure was controlled at 1 bar using the C-rescale barostat ($\tau_p$ = 3 ps). Periodic boundary conditions were applied in all three directions for all simulations.Each system was simulated for 200,000,000 steps, corresponding to a total time of 2.5 μs.[7]

All molecular visualizations were generated with PyMOL 2.5[8]. Clustering analysis was performed by defining a 0.6 nm cutoff distance between neighboring peptides to determine whether they belong to the same aggregate. The time evolution of the number of peptides within the largest aggregate was monitored throughout the simulation (Fig.S1). The structural dynamism of the systems was quantified using residue-wise root mean square fluctuations (RMSF). Analyses were based on the final 1 μs of three independent simulations, including only those systems that had reached equilibrium after the initial 1 μs. The equilibration state was verified by root mean square deviation (RMSD) profiles (Fig.S2) and energy stability analysis (Fig.S4). The solvent-accessible surface area (SASA) indirectly reflects the conformational states and relative positional changes of molecules within the aggregates or at the interfaces (Fig.S2c). The RDF curves reveal the spatial distribution patterns and local structural features of intermolecular arrangements within the systems (Fig. S3).

The peptide structures were first predicted using AlphaFold3 [2], and subsequently coarse-grained into MARTINI models using martinize2 within the Vermouth framework. To better capture the secondary structural characteristics of the sequences, input topology files were manually adjusted during CG mapping, with helical regions explicitly reassigned as $\beta$-sheet types [3].

All FIKVAV-derived peptides—namely FIKVAV, FFIKVAV, FFFIKVAV, and FFFSIKVAV—contain only one charged residue, the naturally protonated lysine (+1). Experimental evidence confirmed that these peptides are capable of spontaneous self-assembly into fibrous aggregates. Therefore, no protonation state modifications were applied during coarse-graining, and the default protonation states defined in the MARTINI force field were retained. As a result, the net charge for each peptide variant was set to +1.

Simulations were conducted in two stages within a cubic simulation box of 21.5 × 21.5 × 21.5 $nm^3$, solvated with MARTINI water, and neutralized with sufficient counterions. In the first stage, 300 peptide molecules were randomly placed in the box with a minimum inter-peptide distance of 3 Å, corresponding to a concentration of 50 mM and a peptide weight fraction ranging from 3.46% to 5.63%, depending on sequence length. This concentration is within the range commonly used for accelerating self-assembly simulations and is up to 10-fold higher than experimental conditions. The systems were equilibrated for 2.5 μs .

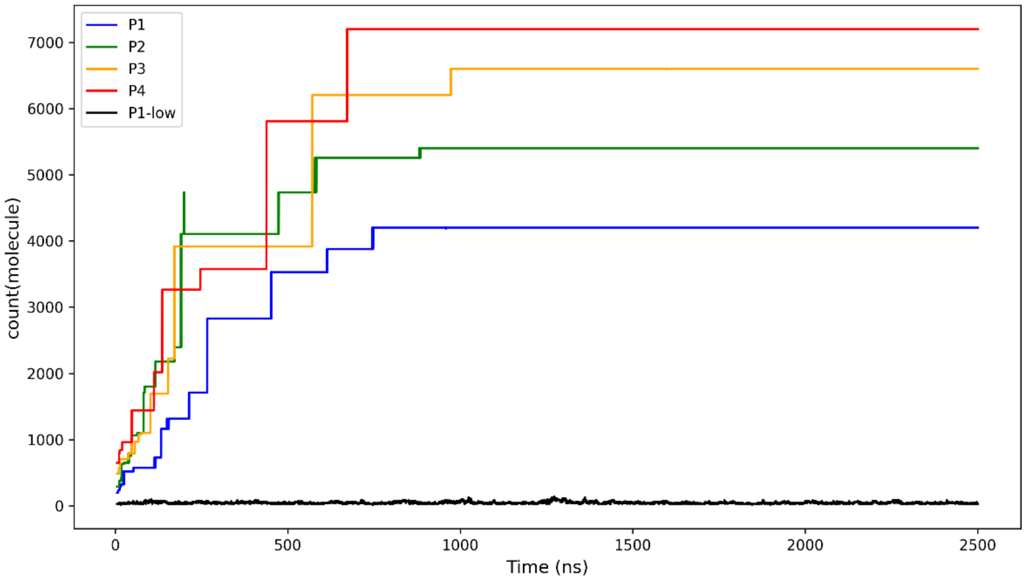


**Figure S1.** Temporal evolution of the number of atoms in peptide aggregates for different systems. **P1**-low denotes the **P1** system at a concentration of 8.3 mM.

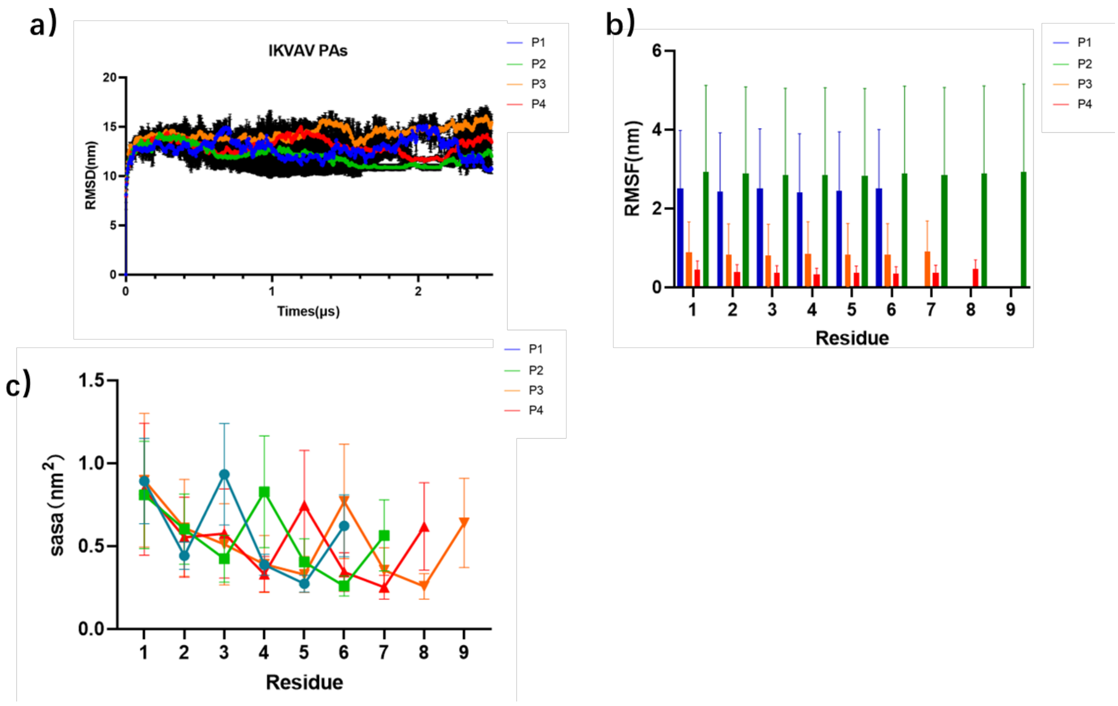


**Figure S2. (a)** Root-mean-square deviation (RMSD) of different peptide systems. **(b)** Residue-wise solvent accessible surface area (SASA) for different peptide systems. **(c)** Residue-wise root-mean-square fluctuation (RMSF) for different peptide systems.

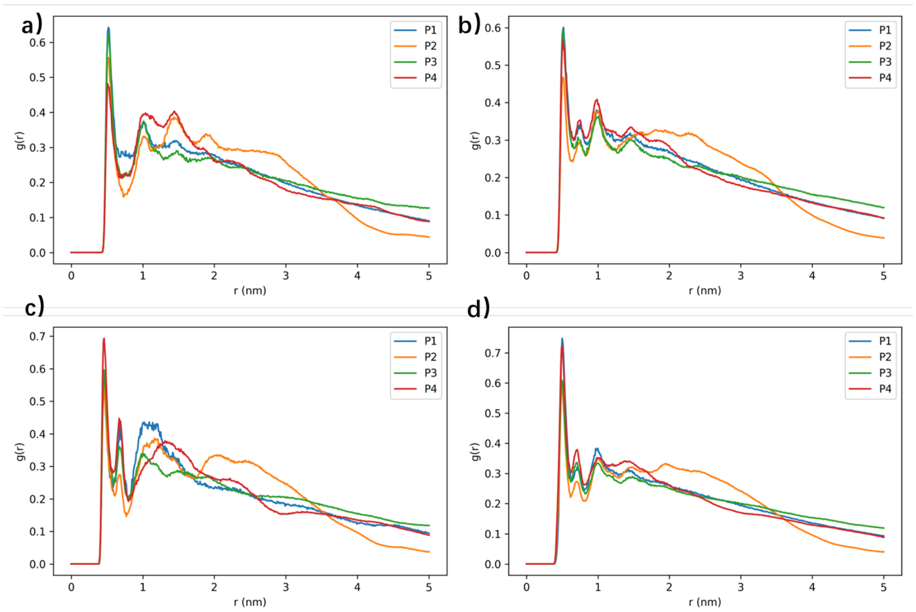


**Figure S3.** Normalized radical distribution function (RDF) curves: **(a)** backbone–backbone, **(b)** backbone–side chain, **(c)** Phe–Phe, and **(d)** side chain–side chain.

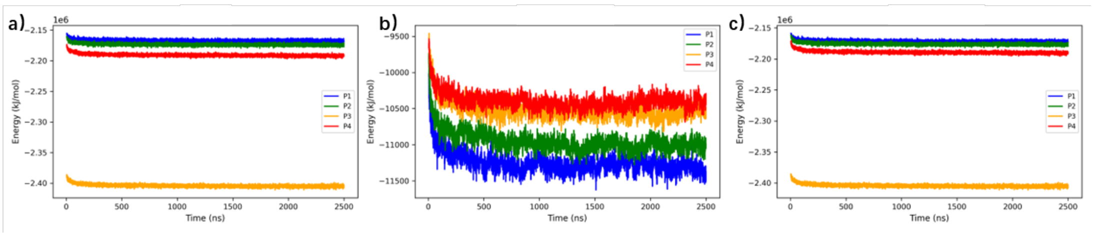


**Figure S4.** Energy analysis of the aggregation process. Panels (a)–(c) correspond to the following energy terms: **(a)** short-range van der Waals (Lennard-Jones, LJ), **(b)** short-range Coulomb interactions, and **(c)** total Potential energy.

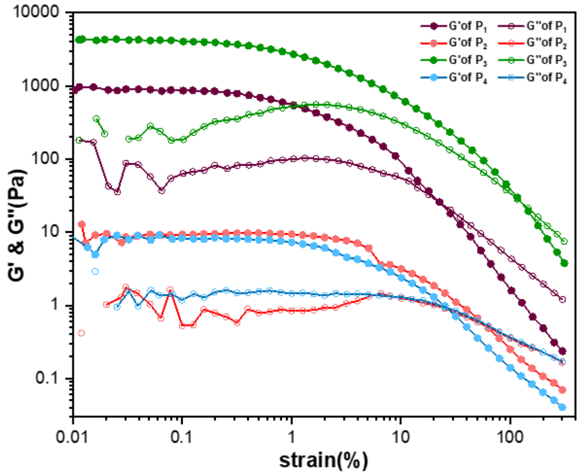


**Figure S5.** Strain amplitude sweep of 4 peptide hydrogels measured at constant frequency.

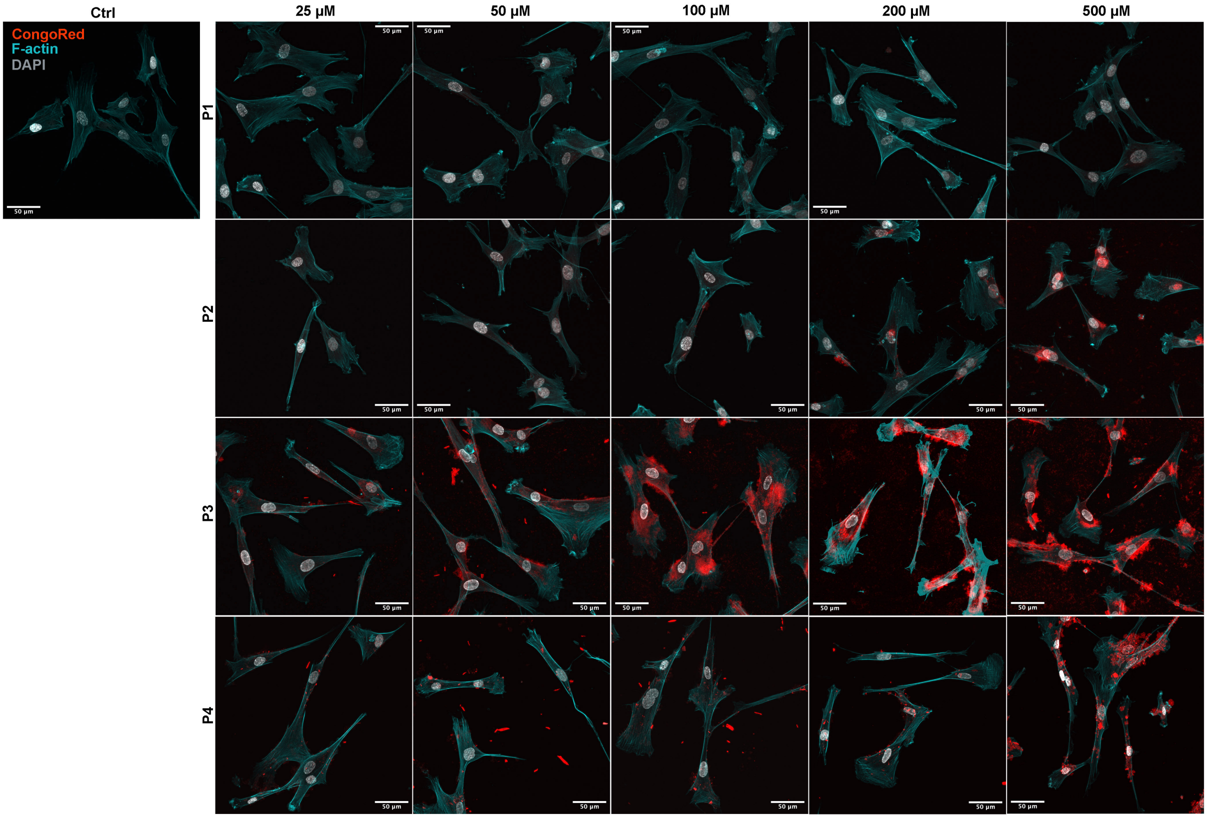


**Figure S6.** Fluorescence images of MSCs with or without peptide treatment at various concentrations for 1 day, co-stained with ActinGreen (cyan), DAPI (gray), and Congo Red (red). The scale bar represents 50 μm.

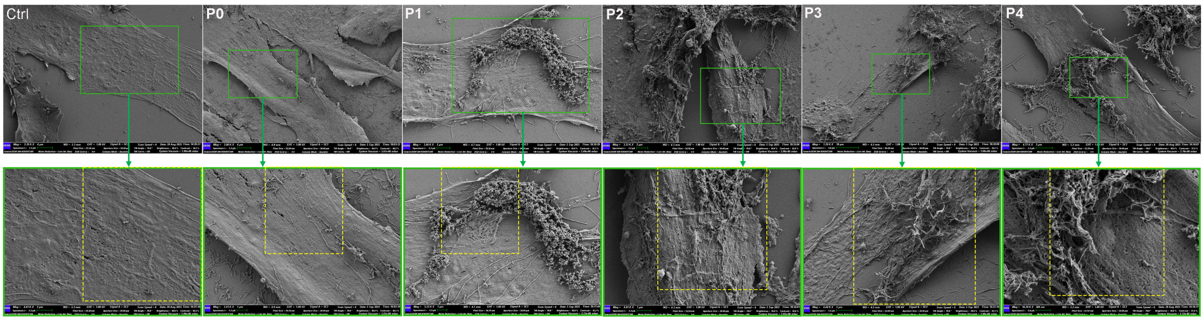


**Figure S7.** Zoomed-out SEM images corresponding to Figure 2b. The regions shown in the enlarged SEM images in Figure 2b are highlighed with yellow dashed-line squares.

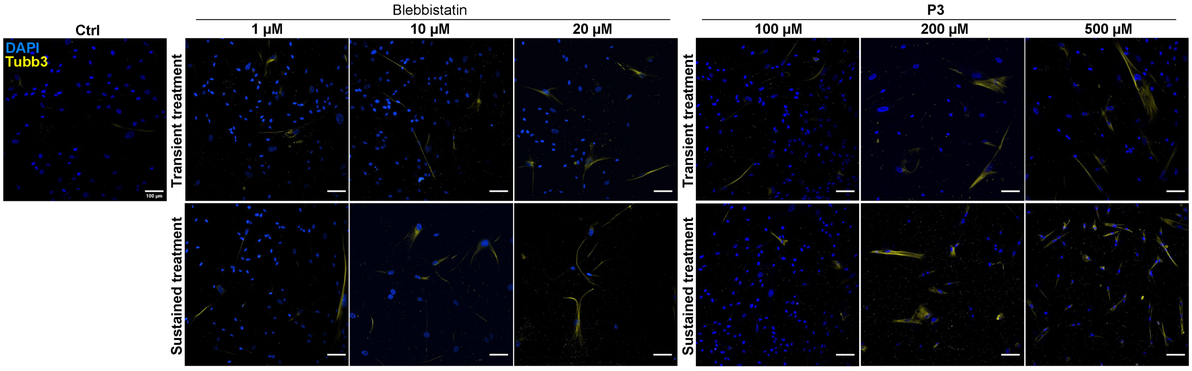


**Figure S8.** Representative immunofluorescence images corresponding to Figure 2c, with DAPI (blue), and TUBB3 (yellow). The scale bars represent 100 µm.

#### Cell Live/Dead Assay

Cells were seeded in a 96-well flat-bottom tissue culture plate (Merck, Germany; CLS3596) at a density of 100 cells per well and incubated overnight to allow cell adhesion. The medium was then replaced with fresh medium containing 200 μM of each peptide compound and 10 μM Blebbistatin (MCE, HY13441). Cells were cultured for an additional 14 Days, and the medium was renewed every 3 days with freshly prepared solutions containing the same concentrations of peptides and small molecules as at the start of the experiment. After treatment, the culture medium was gently aspirated, and cells were washed twice with PBS to remove residual serum. Live and dead cells were then stained using calcein-AM and propidium iodide (PI) from the Double Staining kit (Beyotime, China; C2015L) according to the manufacturer's protocol. Fluorescence microscopy was acquired using a Nikon ECLIPSE Ts2 microscope.

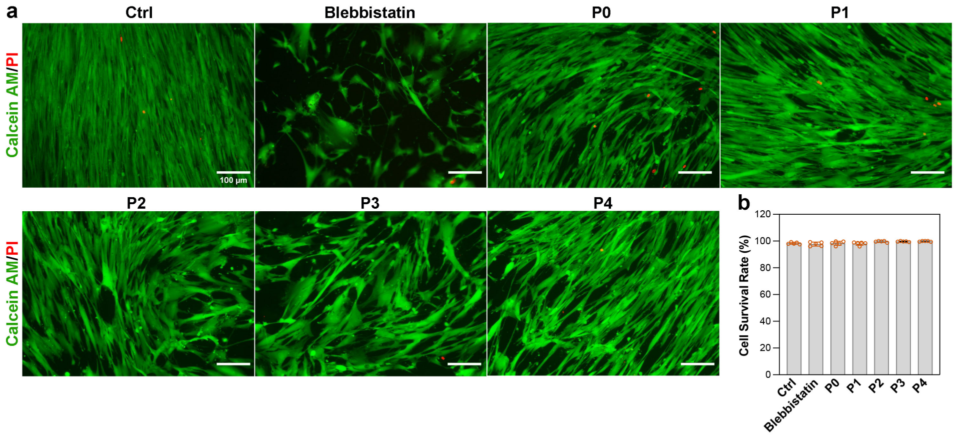


**Figure S9. (a)** Fluorescence images of BMSCs cultured under various conditions for 14 days, stained with Calcein AM/PI Double Staining Kit. Scale bars represent 100 μm. **(b)** Quantitative analysis of cell survival rate.

#### RNA Isolation and Quantitative PCR (qPCR) Analysis

**Table S1.** The primers used for real-time PCR.

| mRNA | Description | Oligonucleotide |
|---|---|---|
| MAP2 | Microtubule-Associated Protein 2 | Forward 5’- TGGTGCCGAGTGAGAAGAAG-3’<br>Reverse 5’- AGTGGTTGGTTAATAAGCCGAAG-3’ |
| TUBB3 | Tubulin Beta 3 Class III | Forward 5’- CCGAAGCCAGCAGTGTCTAA-3’<br>Reverse 5’- AGGCCTGGAGCTGCAATAAG-3’ |
| NEFH | Neurofilament Heavy Chain | Forward 5’-CCGTCATCAGGCCGACATT -3’<br>Reverse 5’-GTTTTCTGTAAGCGGCTATCTCT -3’ |
| RPLP0 | Ribosomal Protein, Large Subunit, P0 | Forward 5’- CGTCCTCGTGGAAGTGACAT -3’<br>Reverse 5’- ATCTGCTTGGAGCCCACATT -3’ |

**Calcium-Flux Imaging**

Supporting Video cllip1

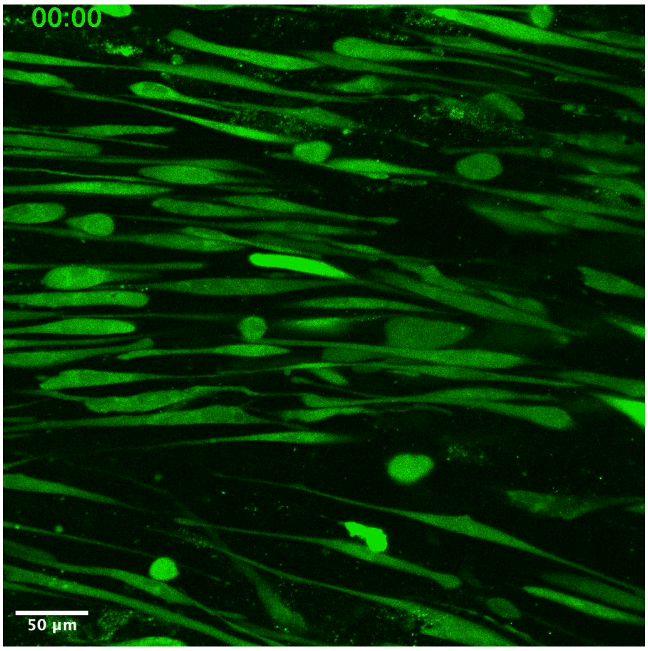

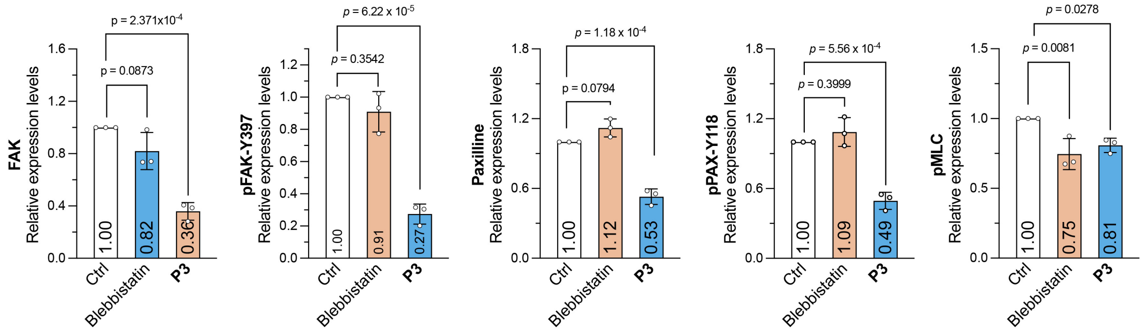


**Figure S10.**Quantitative analysis of relative protein expression levels examined via western blotting. Three independent experiments were performed.

#### References and Notes


[1] Xunwu Hu, Sona Rani Roy, Chengzhi Jin, Guanying Li, Qizheng Zhang, Natsuko Asano, Shunsuke Asahina, Tomoko Kajiwara, Atsushi Takahara, Bolu Feng, Kazuhiro Aoki, Cehnjie Xu, Ye Zhang* "Control Cell Migration by Engineering Integrin Ligand Assembly" *Nature Communications*, 2022, 13:5002.

[2] Abramson, J., Adler, J., Dunger, J. et al. "Accurate structure prediction of biomolecular interactions with AlphaFold 3." *Nature*, 2024, 630, 493–500.

[3] Kroon Peter C, Grunewald Fabian, Barnoud Jonathan, van Tilburg Marco, Brasnett Chris, de Souza Paulo Cesar Telles, Wassenaar Tsjerk A, Marrink Siewert-Jan J. "Martinize2 and Vermouth: Unified Framework for Topology Generation." eLife, 2023, 12:RP90627

[4] Álvarez, Z., Kolberg‐Edelbrock, A. N., Sasselli, I. R., Ortega, J. A., Qiu, R., Syrgiannis, Z., Mirau, P. A., Chen, F., Chin, S. M., Weigand, S., Kiskinis, E., & Stupp, S. I. "Bioactive scaffolds with enhanced supramolecular motion promote recovery from spinal cord injury." Science, 2021, 374, 848–856.

[5] Bussi, G., & Parrinello, M. "Accurate sampling using Langevin dynamics with stochastic cell rescaling." Physical Review E, 2007, 75, 056707.

**Attachment 1.** LC-MS spectra of peptide **P0** (IKVAV).

Column : 4.6mm*250mm, SinoChrom ODS-BP
Solvent A : 0.1% trifluoroacetic in 100% acetonitrile
Solvent B : 0.1% trifluoroacetic in 100% water

| Gradient : | A | B |
|---|---|---|
| 0.01min | 13% | 87% |
| 25min | 38% | 62% |
| 25.1min | 100% | 0% |
| 30.0min | STOP | |

Flow rate : 1.0ml/min
Wavelength : 220nm
Volume : 5ul

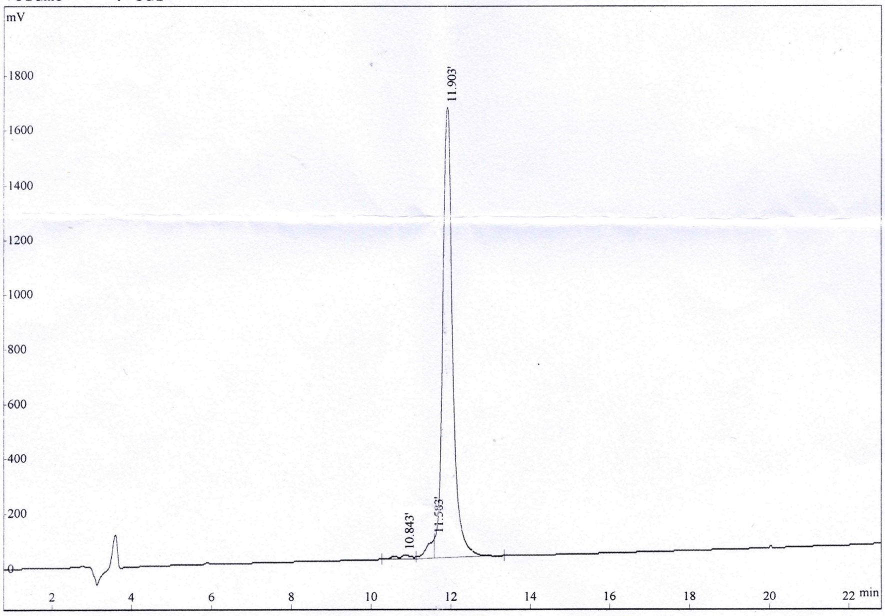


| Rank | Time | Conc. | Area | Height |
|---|---|---|---|---|
| 1 | 10.843 | 1.033 | 294510 | 12289 |
| 2 | 11.583 | 3.043 | 867281 | 67701 |
| 3 | 11.903 | 95.92 | 27338861 | 1640315 |
| Total | | 100 | 28500652 | 1720305 |

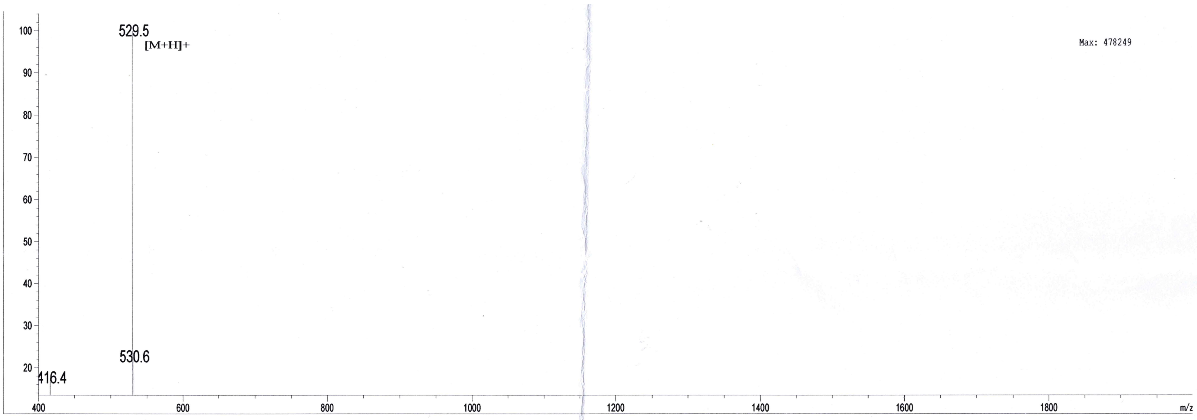


| Sample Description | | Instrument | Agilent-6125B | | |
|---|---|---|---|---|---|
| Analyzed date: | 2025-06-26 | Probe: | ESI | Probe Bias: | +4.5kv |
| Analyst: | YU | Nebulizer Gas Flow: | 1.5L/min | Detector: | 1.5kv |
| Sample: | 定制肽 3 IV-5 | CDL: | -20.0v | T. Flow: | 0.2ml/min |
| M.W.: | 528.69 | CDL Temp.: | 250 °C | B. Conc.: | 50%H2O/50%ACN |
| Lot. No.: | P250619-LC061697 | Block Temp.: | 200 °C | | |

**Attachment 2.** $^{1}$H NMR spectrum of peptide **P0** (IKVAV).

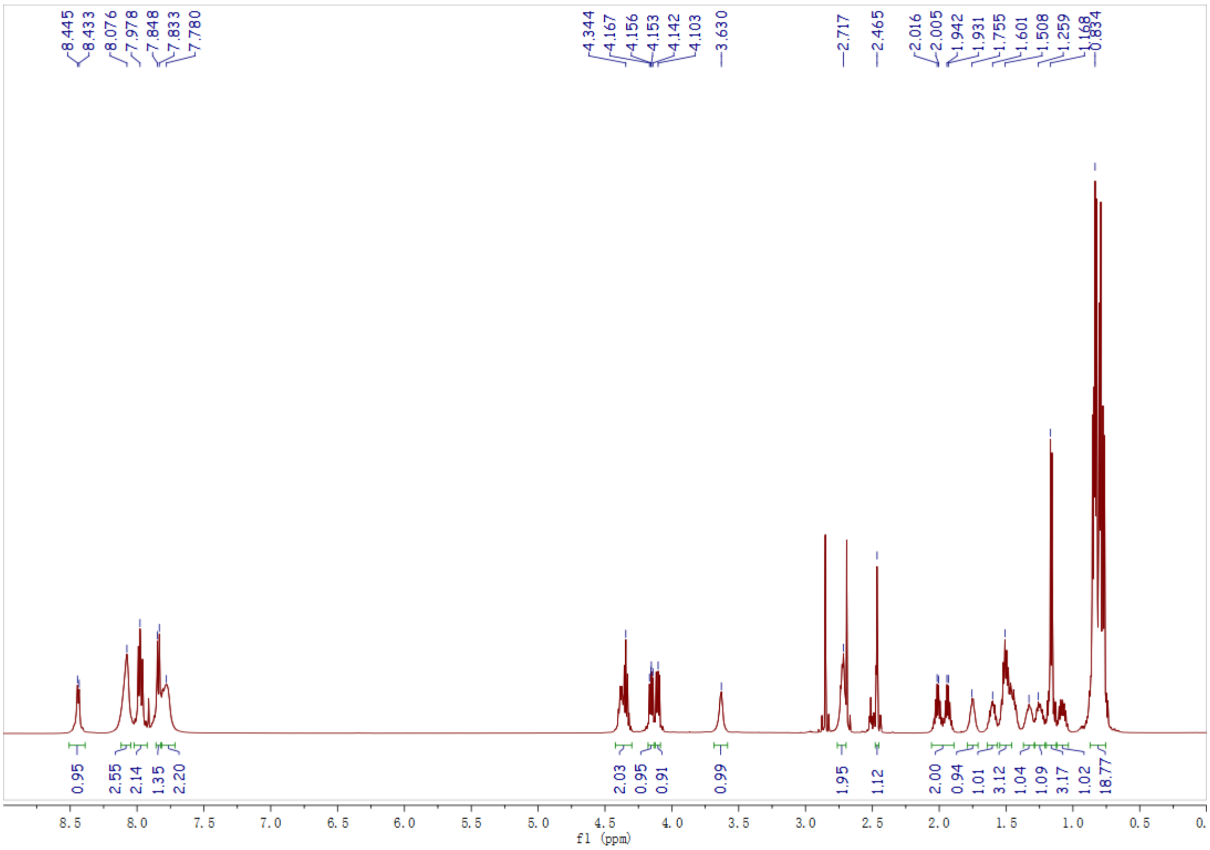

**Attachment 3.** LC-MS spectra of peptide **P1** (FIKVAV).

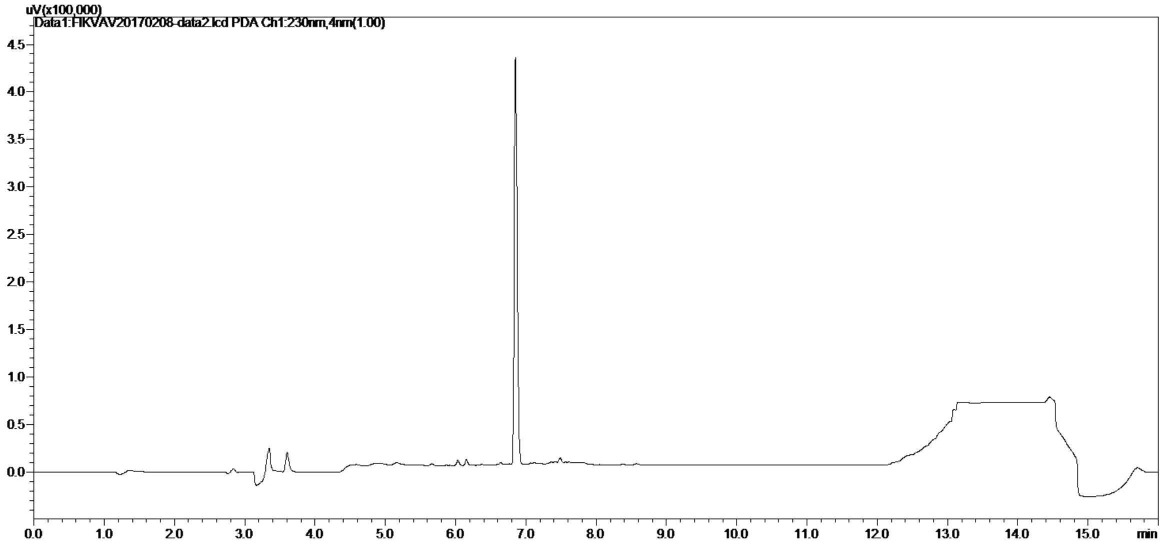


D:\Roy_BRS\...\FIKVAV20170208-data3 2/15/2017 2:44:27 PM

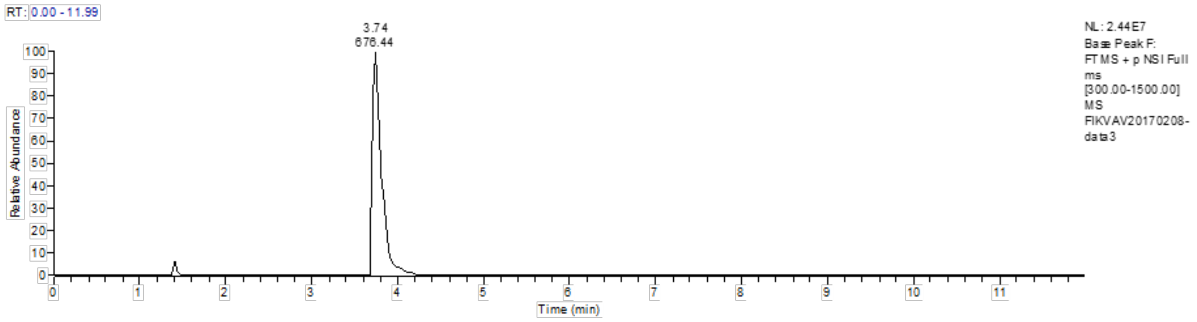


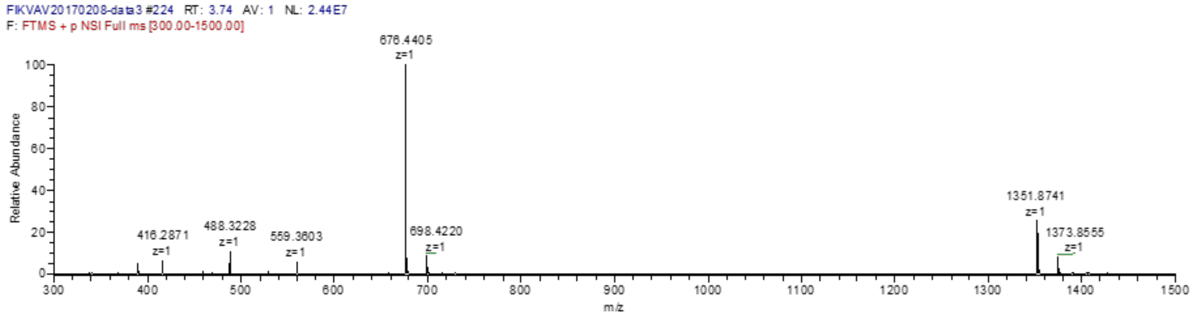

**Attachment 4.** [1]H NMR spectrum of peptide **P1** (FIKVAV).

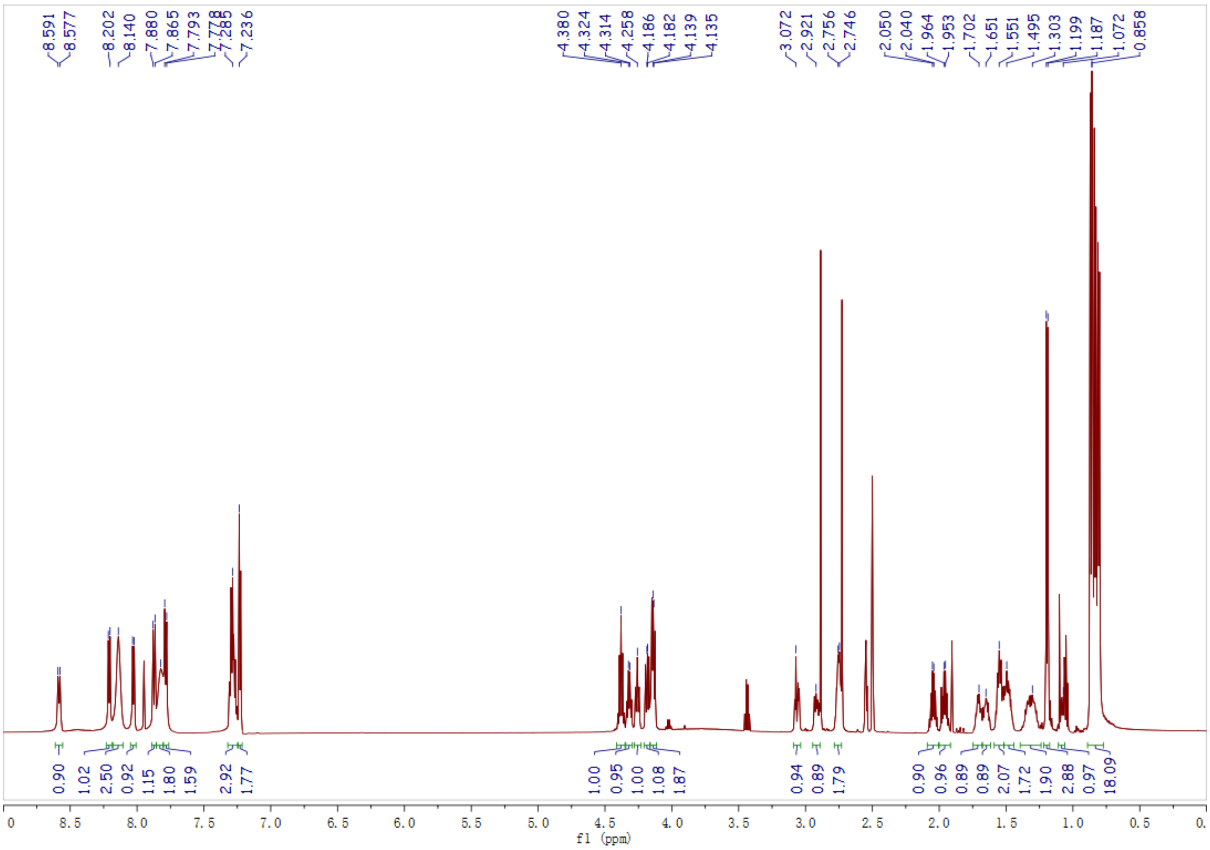

**Attachment 5.** LC and HRMS spectra of peptide **P2** (FFIKVAV).

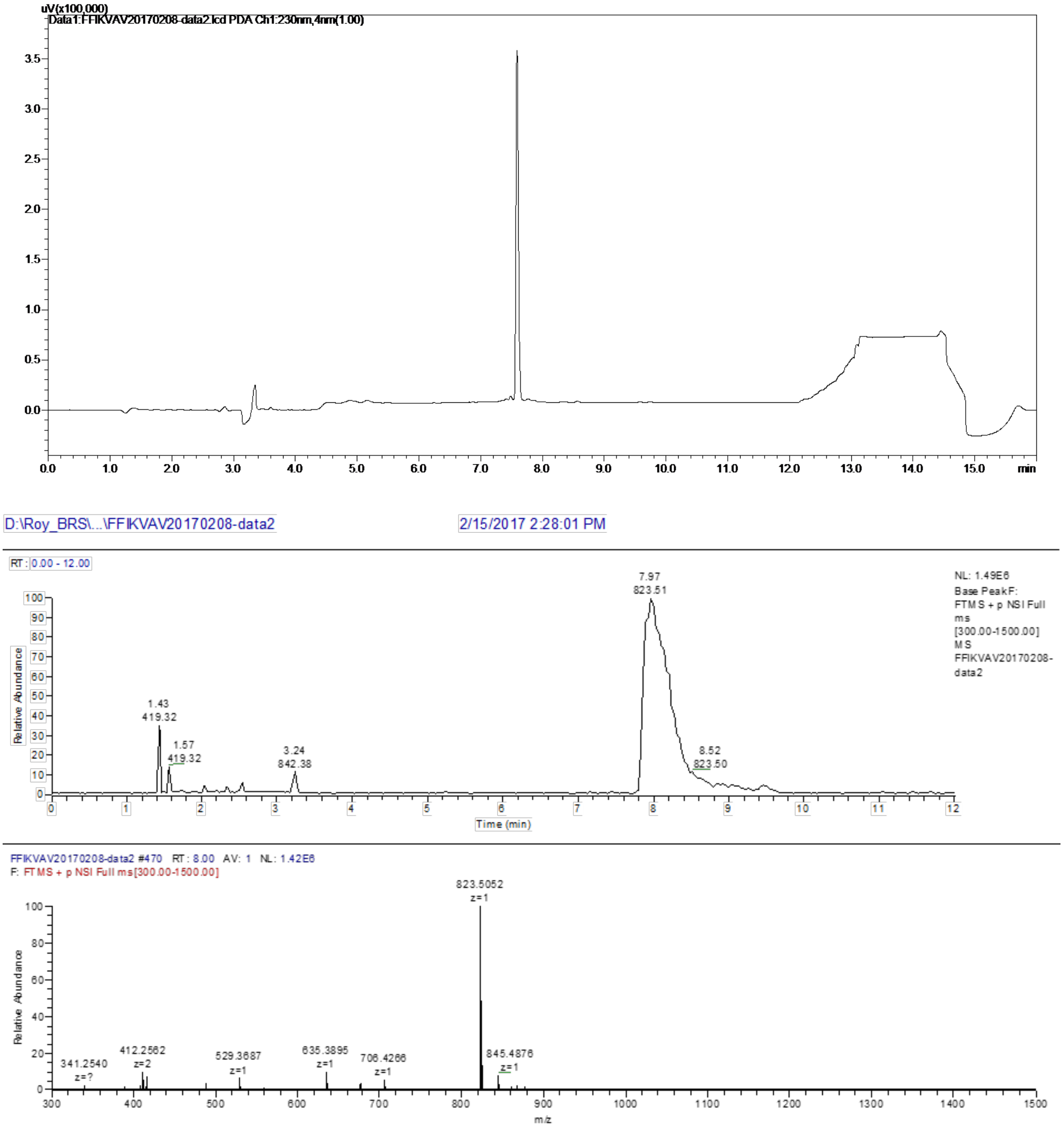

**Attachment 6.** $^{1}$H NMR spectrum of peptide **P2** (FFIKVAV).

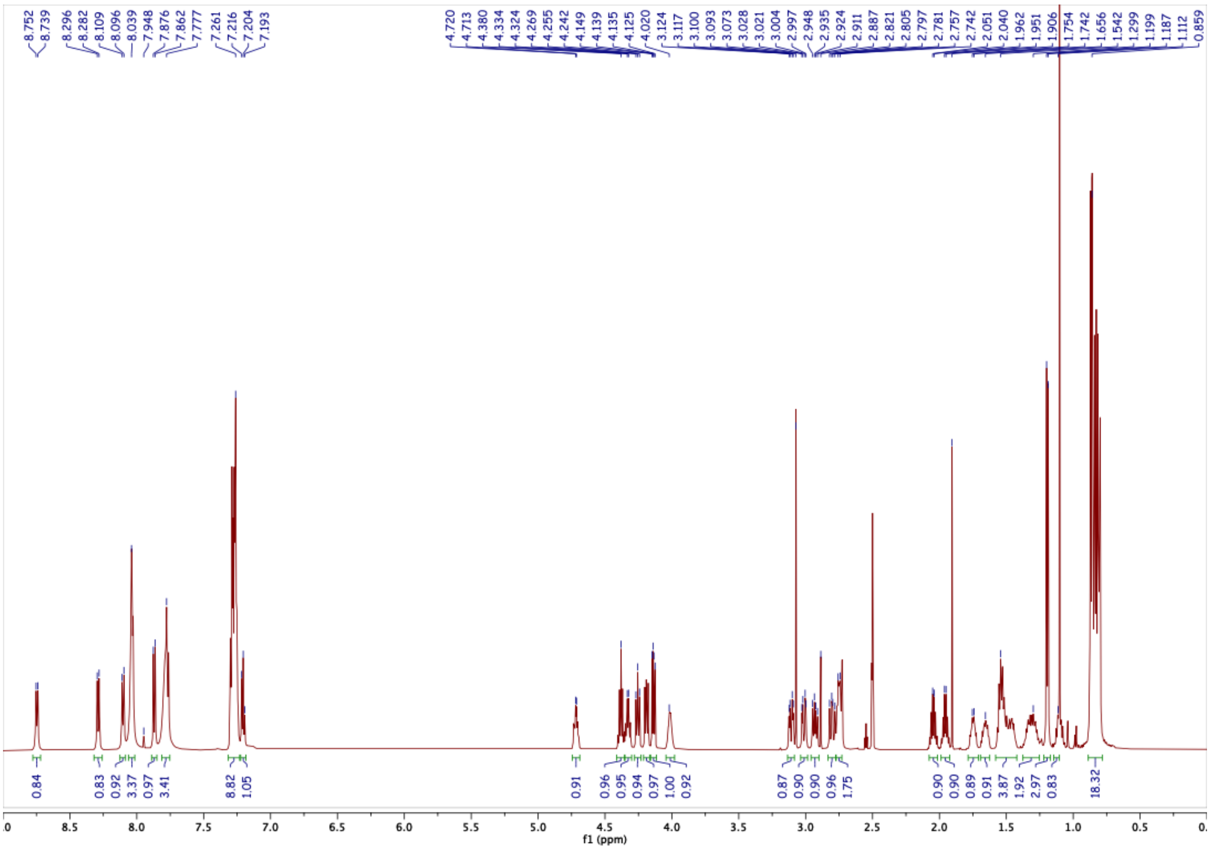

**Attachment 7.** HPLC and HRMS spectra of peptide **P4** (FFFSIKVAV).

| | | | |
|---|---|---|---|
| Column | : Shim-pack GIST C18 5 μm 4.6 I.D. x 250 mm | | |
| Solvent A | : 0.1% trifluoroacetic acid in water | | |
| Solvent B | : 0.1% trifluoroacetic acid in acetonitrile | | |
| Gradient | : Time (min) | Pump A (%) | Pump B (%) |
| | 0.00 | 70.0 | 30.0 |
| | 25.00 | 30.0 | 60.0 |
| | 26.00 | 0.0 | 100.0 |
| | 30.00 | 0.0 | 100.0 |
| | 31.00 | 70.0 | 30.0 |
| | 35.00 | 70.0 | 30.0 |
| Flow rate | : 1.0 mL/min | | |
| Wavelength | : 220 nm | | |
| Volume | : 30 μL | | |

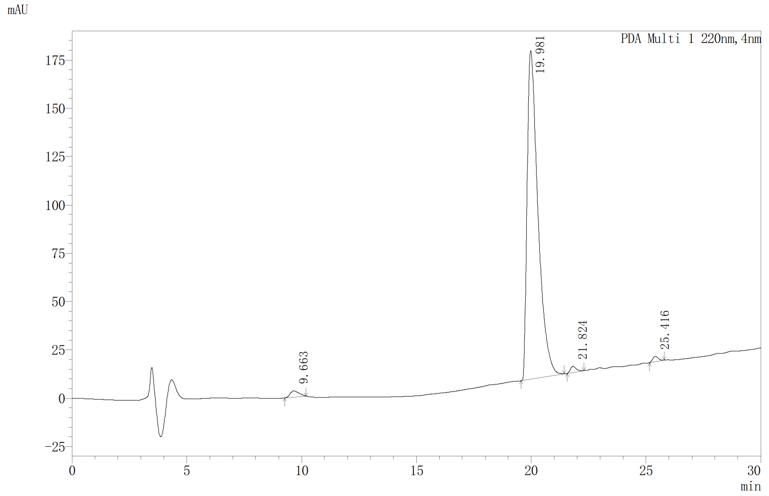


| Rank | Time | Area | Height | Area% |
|---|---|---|---|---|
| 1 | 9.663 | 86725 | 3237 | 1.501 |
| 2 | 19.981 | 5591239 | 169885 | 96.752 |
| 3 | 21.824 | 54651 | 3079 | 0.946 |
| 4 | 25.416 | 46315 | 2772 | 0.801 |
| Total | | 5778930 | 178974 | 100.000 |

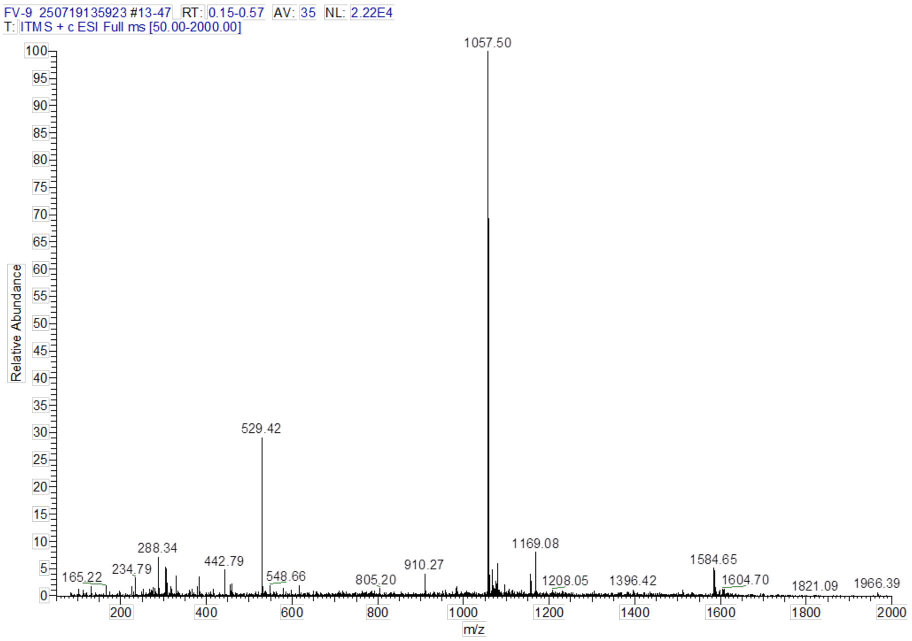

**Attachment 8.** [1]H NMR spectrum of peptide **P4** (FFFSIKVAV).

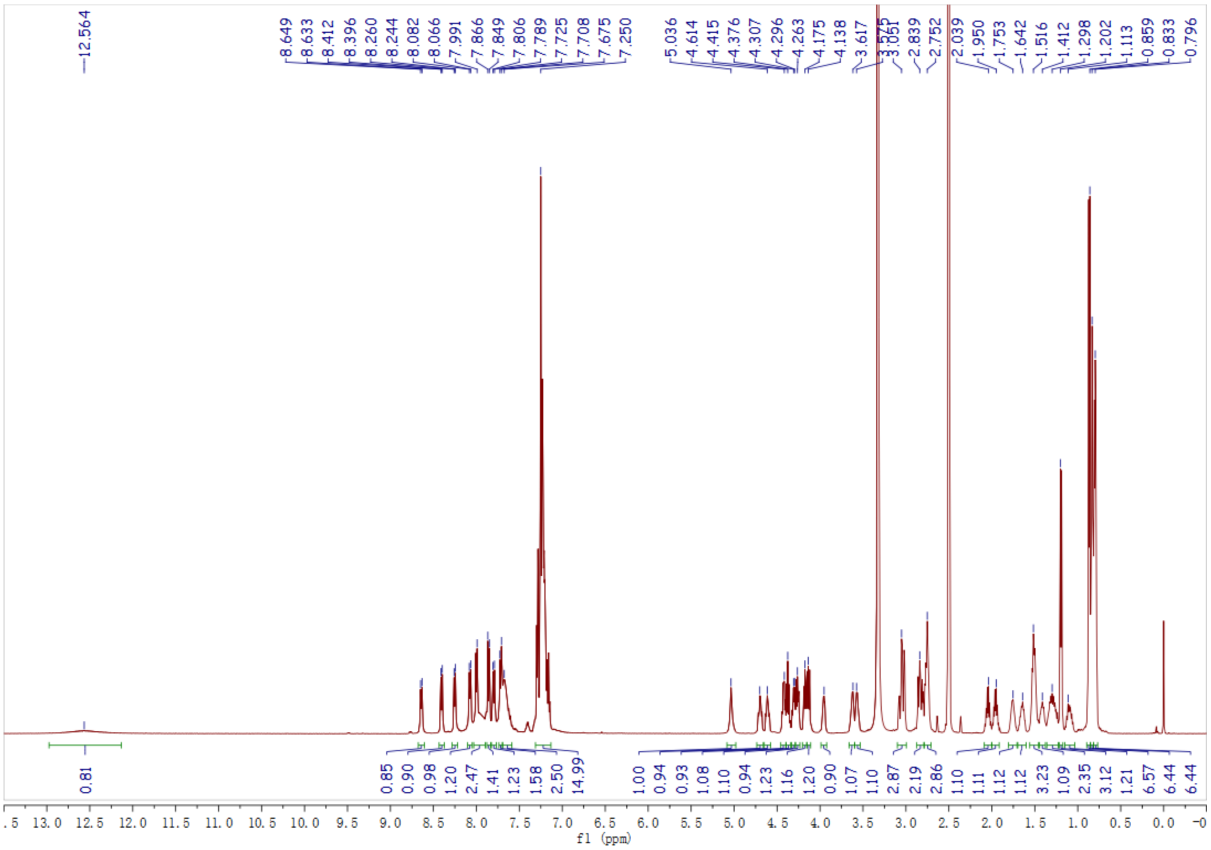
